# The ground offers acoustic efficiency gains for crickets and other calling animals

**DOI:** 10.1101/2022.11.13.516353

**Authors:** Erin E. Brandt, Sarah Duke, Honglin Wang, Natasha Mhatre

## Abstract

Male crickets attract females by producing calls with their forewings. Louder calls travel further and are more effective at attracting mates. However, crickets are much smaller than the wavelength of their call, and this limits their power output. A small group called tree crickets make acoustic tools called baffles which reduce acoustic short-circuiting, a source of dipole inefficiency. Here, we ask why baffling is uncommon among crickets. We hypothesize that baffling may be rare, because like other tools they offer insufficient advantage for most species. To test this, we modelled the calling efficiencies of crickets within the full space of possible natural wing sizes and call frequencies, in multiple acoustic environments. We then generated efficiency landscapes, within which we plotted 112 cricket species across 7 phylogenetic clades. We found that all sampled crickets, in all conditions, could gain efficiency from tool use. Surprisingly, we also found that calling from the ground significantly increased efficiency, with or without a baffle, by as much as an order of magnitude. We found that the ground provides some reduction of acoustic short-circuiting but also halves the air volume within which sound is radiated. It simultaneously reflects sound upwards, allowing recapture of a significant amount of acoustic energy through constructive interference. Thus, using the ground as a reflective baffle is an effective strategy for increasing calling efficiency. Indeed, theory suggests that this increase in efficiency is accessible not just to crickets, but to all acoustically communicating animals whether they are dipole or monopole sound sources.

**Significance Statement:** Loudness is a crucial feature in acoustic communication. Animals attracting mates or warding off predators are expected to make themselves as loud as possible. Two long-standing, seemingly unrelated unsolved problems regarding loudness in the field of animal communication are: the rarity of acoustic tool use, and animals that call from reflective ground-like surfaces, known to be an impediment to sound propagation. These two ideas are related; by refocusing analysis from sound propagation to sound radiation, we show that the ground is not an impediment, but rather an acoustic aid that can boost loudness more than tool use. We also show that calling from a reflective surface is an alternative strategy to maximize call loudness that is available to all animals.

## Introduction

Male crickets make loud advertisement calls to attract females who use these calls to locate mates (1). Louder calls travel further, cover more area, and attract more females (2–4). When faced with a choice, females prefer louder calls (2, 5). Being louder therefore has implications for mating success and evolutionary fitness in these singing insects. However, despite the apparent loudness of a nighttime chorus, cricket calls are acoustically constrained by a phenomenon known as ‘acoustic short-circuiting’ specific to dipole sound sources (6, 7). Cricket wings are sound radiators that vibrate back and forth in the air like pistons. As a wing moves in one direction, the air in front of the wing is compressed, and the air behind it is rarified. These two changes in pressure travel away as waves as the motion is periodically repeated. However, the waves on either side of the wing are of opposite phase and interfere destructively where they meet, at the edges of the wing. Thus, less sound is radiated, reducing sound radiation efficiency (6, 8). The smaller the wings of a cricket with respect to the wavelength of the sound it makes, the higher the short-circuiting and associated loss of efficiency. Indeed, the few crickets that have been studied are small and experience significant short-circuiting (9).

To compensate for the power lost to acoustic short-circuiting, a few tree cricket species build and use an acoustic tool known as a baffle (5–7, 10). A baffle consists of a leaf with a hole chewed by the cricket near the middle of the leaf. When the size of the leaf and hole are optimal, such structures reduce acoustic short-circuiting and increase efficiency by as much as 10 dB compared to unbaffled calling, reflecting a tripling of sound pressure levels (7). However, despite their benefits, only a handful of species among thousands make baffles, all within the sub-family Oecanthinae (5, 7, 10, 11).

Given the obvious benefits, why is acoustic baffle use rare in crickets? Tree cricket baffles are tools, and tool use is generally rare (12–14). Indeed, few species use tools, whether crickets, other invertebrates or even vertebrates. Invertebrate tool use, however, seems especially rare. For example, 56 independent occurrences of tool use were found in mammals, whereas only 13 occurrences were found in the significantly more speciose insects (14). Two hypotheses from the tool use literature, the “cognitive capacity” and the “lack of utility” hypotheses offer two different reasons for this rarity. The “cognitive capacity” hypothesis suggests that complex tool use behaviors are less likely to evolve in animals with smaller brains and lower cognitive capacity. This is an unlikely explanation since many animals with relatively low cognitive capacities do use, and even make, tools which themselves are not necessarily complex objects. Small-brained animals are even known to make very complex and highly functionally optimized habitation structures which do not require high cognitive capacity (7, 15).

A competing hypothesis is the “lack of utility” hypothesis which posits that tool behavior can evolve regardless of cognitive capacity, but that its evolution requires an ecological context in which it confers sufficient selective advantage (15). Only species that can achieve higher gains from tool use than from other strategies (e.g., morphological features, site selection) are likely to evolve tool using behavior.

To test the lack of utility hypothesis, we must quantify tool utility and use of the tool must have implications for evolutionary fitness. It is often difficult to meet these two conditions. Work in chimpanzees has directly quantified tool utility by evaluating how much caloric value can be gained by using a tool to exploit an otherwise unexploitable resource (16). Other studies have made more indirect arguments; work in sea otters has shown that tools are employed more frequently in populations in which tough prey require tools to access them (17). In capuchin monkeys, larger individuals who can more effectively use tools to crack nuts are more likely to use tools (18). However, few studies quantitatively assess the lack of utility hypothesis, particularly outside the context of food.

Baffle use in crickets is an ideal system in which to test the lack of utility hypothesis. First, baffle use is rare and second, we can directly measure its acoustic utility in terms of an increase in sound radiation efficiency (12). Finally, baffle use has been shown to have real fitness implications, by increasing the number of mates attracted to a given male, and also by increase mating duration, both processes likely to increase reproductive success (2).

Therefore, in this study, we tested the lack of utility hypothesis across a large group of singing insects, the true crickets or Grylloidea. We used the finite element method (7, 19, 20) to quantify baffle utility in two ways. First, we ascertained the range of sizes of the sound radiator (cricket wings) and frequency ranges of the calls used by 112 crickets, spread over the cricket phylogeny and used it to define an acoustic-morphospace (Fig 1). Then we quantified a volumetric sound radiation efficiency (SRE_V_), averaged across all positions in this space, specifically the relation between the radiator vibration velocity amplitude (m/s), and the sound pressure levels (Pa) that are generated, similar to the previous study on tree cricket baffle efficiency (7). Therefore, the dimensions of SRE_V_ would be Pa/(m/s). Since the amplitude of the radiator vibration velocity would reflect the effort applied by the animal, this measure of efficiency captures a significant component of the relationship between calling effort and output. By plotting SRE_V_ over the complete acoustic-morphospace, we were able to generate efficiency landscapes. This landscape allowed us to ascertain the efficiencies of both unbaffled and baffled crickets and enabled us to fully investigate all possible crickets, even those that did not appear in our sample.

**Figure 1.**
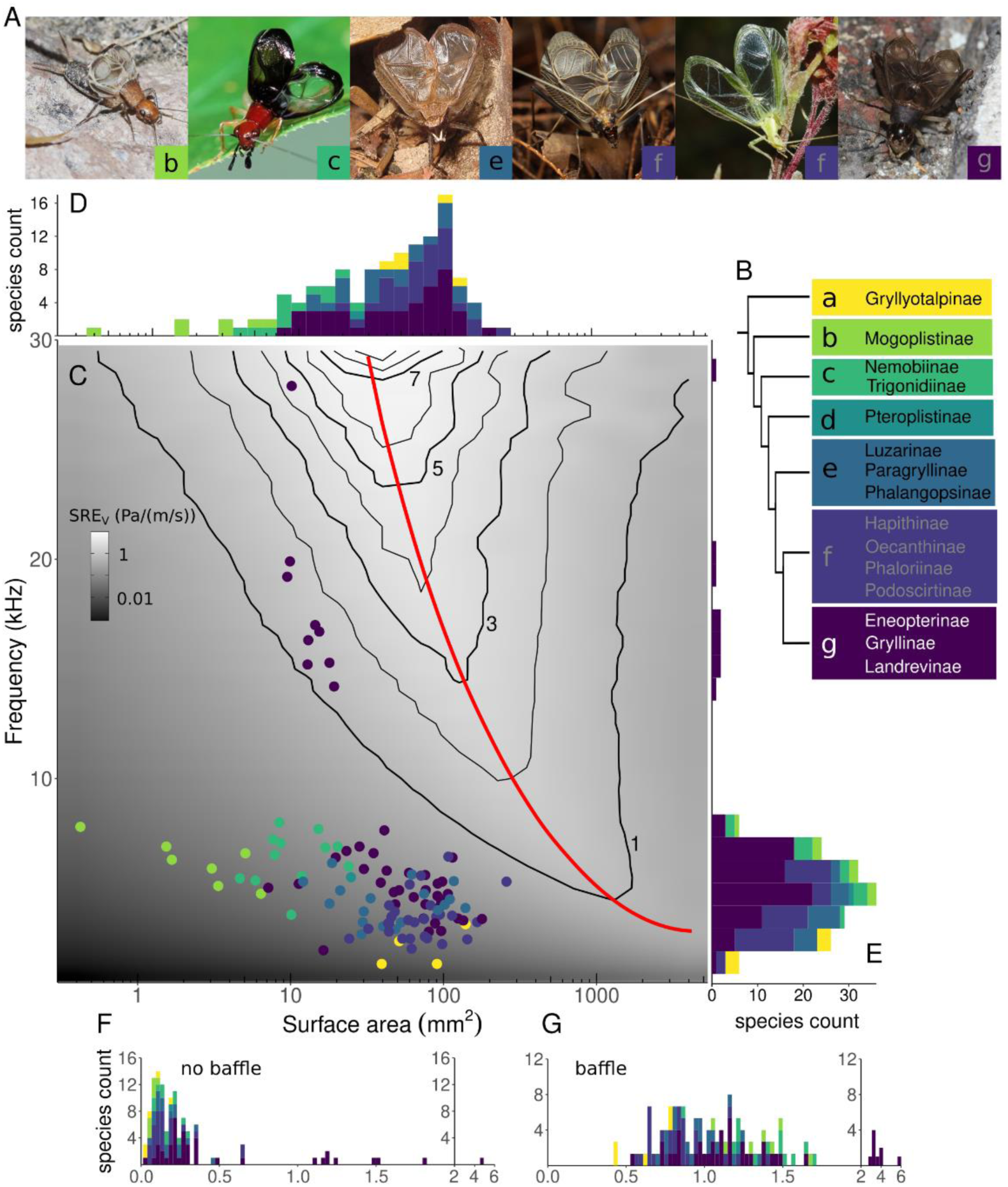
All crickets could increase efficiency by baffled calling. Sound radiation efficiency (SRE) landscape across the acoustic-morphospace of crickets. **A.** Representative images of cricket males with wings raised in calling posture. Species pictured are (from left): *Hoplosphyrum boreale*, *Phyllopalpus pulchellus, Lerneca inalata, Meloimorpha japonica, Oecanthus quadripunctatus*, and *Turanogryllus eous.* (Photos from left: James Bailey, Wilbur Hershberger, Richard C. Hoyer, Ryosuke Kuwahara, James P. Bailey and Taewoo Kim). Boxes with letter indicate the clade of each species. **B.** Phylogeny illustrating the seven clades defined by Chintauan-Marquier et al (2016) and subfamilies within each clade (branch lengths not to scale). **C.** SRE_V_ achieved with different combinations of wing sizes vibrating at different frequencies. Within this space of possibilities, wing areas and call frequencies of all sampled animals are overlaid as points on the SRE landscape. This SRE_V_ is calculated from finite element models. Red line indicates the point for optimal radiator size at each frequency, which corresponds to the radiator size at each frequency that will have minimal short-circuiting and maximal efficiency. A cricket singing from an optimal baffle would also experience minimal short-circuiting and lie on this line, so the place where each animal’s frequency intersects with the line is its baffled SRE_V_. **D, E.** Distributions of calling song frequency and wing size of different animals. Histograms include additional species for which only wing or call measurements were available. **F.** SRE_V_ of each species without use of a baffle. **G.** SRE_V_ of each species with use of an optimal baffle.

In this efficiency landscape, while we are considering a morphological parameter (radiator size), we are not positing that this parameter will be optimized. In biology, trait optimization can be treated as a hypothesis, and can be productive if tested. In fact in some rare instances, it may be found to be true, as was shown by Barlow in the case of diffraction-limited ommatidial-lens sizes in insect eyes (21, 22). Here, we quantitatively define calling efficiency and test if there are alternative paths to higher efficiency such as through tool-use behavior, or through the use of environmental features.

Therefore, to address the complexity of the natural environmental conditions in which crickets call, such as, from close to the ground, and from within vegetation, we generated additional efficiency landscapes. In both these cases, the environment interacts with the sound radiator and the sound emanating from it across spatial scales, and may effectively add or remove any gains from baffling (25, 26). To capture spatial effects, we generated a second metric sound radiation efficiency, this time measured along a transect, (SRE_T_). Here we used the boundary element method and generated new efficiency landscapes which quantified the effects of interacting surfaces on acoustic efficiency as sound propagates away from the caller, under a range of environmental conditions.

Crucially, we considered calling from the ground quite carefully, in terms of its effect on efficiency. In the animal communication literature, typically the ground is considered during propagation and not in relation of sound radiator efficiency. In this context, it is typically seen as a severe impediment to sound propagation by causing significant excess attenuation (27–32). Researchers also focus on the ground effect as degrading temporal structure (31) and directional information (32, 33). While we cannot address temporal structure in our examination of sound radiation efficiency, excess attenuation is accounted for. Additionally, using the same models, we also quantified directionality to test how efficiency might trade off with this biologically crucial feature.

Using these data, we asked whether the rarity of baffle use in crickets is explained by the lack of utility hypothesis. We examined the differences between baffled and unbaffled calling in different realistic scenarios. We expect that known baffle-users will be animals who benefit most from baffle use, and non baffle-users might not accrue as many benefits due to acoustic, morphological or environmental constraints.

## Results

### All crickets would benefit from baffle use in idealized conditions

To capture the natural range of wing sizes and calling frequencies among true crickets, we collected wing surface area and call frequency data for 112 cricket species from a large range of sources (Fig 1, Tables S1, S2, Fig S1). Species were distributed across 7 clades as described by the most recent phylogeny of the Grylloidae or “true cricket” superfamily (34) (Fig 1, Fig S1).

We then constructed finite element models which predicted the sound fields produced by wings of different sizes at different call frequencies for 1086 different combinations which encompassed all the observed frequencies and wing sizes, i.e. the full acoustic-morphospace (Fig 2). In all conditions, wings were modelled as suspended in free space, vibrating with a uniform velocity perpendicular to the wing plane (Fig S2). The model predicts the resulting sound field (see Supplementary methods for details). We then calculate sound radiation efficiency (SRE_V_ (Pa/(m/s))), by taking a volumetric average of the sound pressure level generated (Pa), over a sphere of radius 20 cm around the wing, divided by the time-space average of the wing vibration velocity (m/s). This normalized measure of efficiency enables comparison between species, no matter their actual wing velocity or sound pressure level.

**Figure 2.**
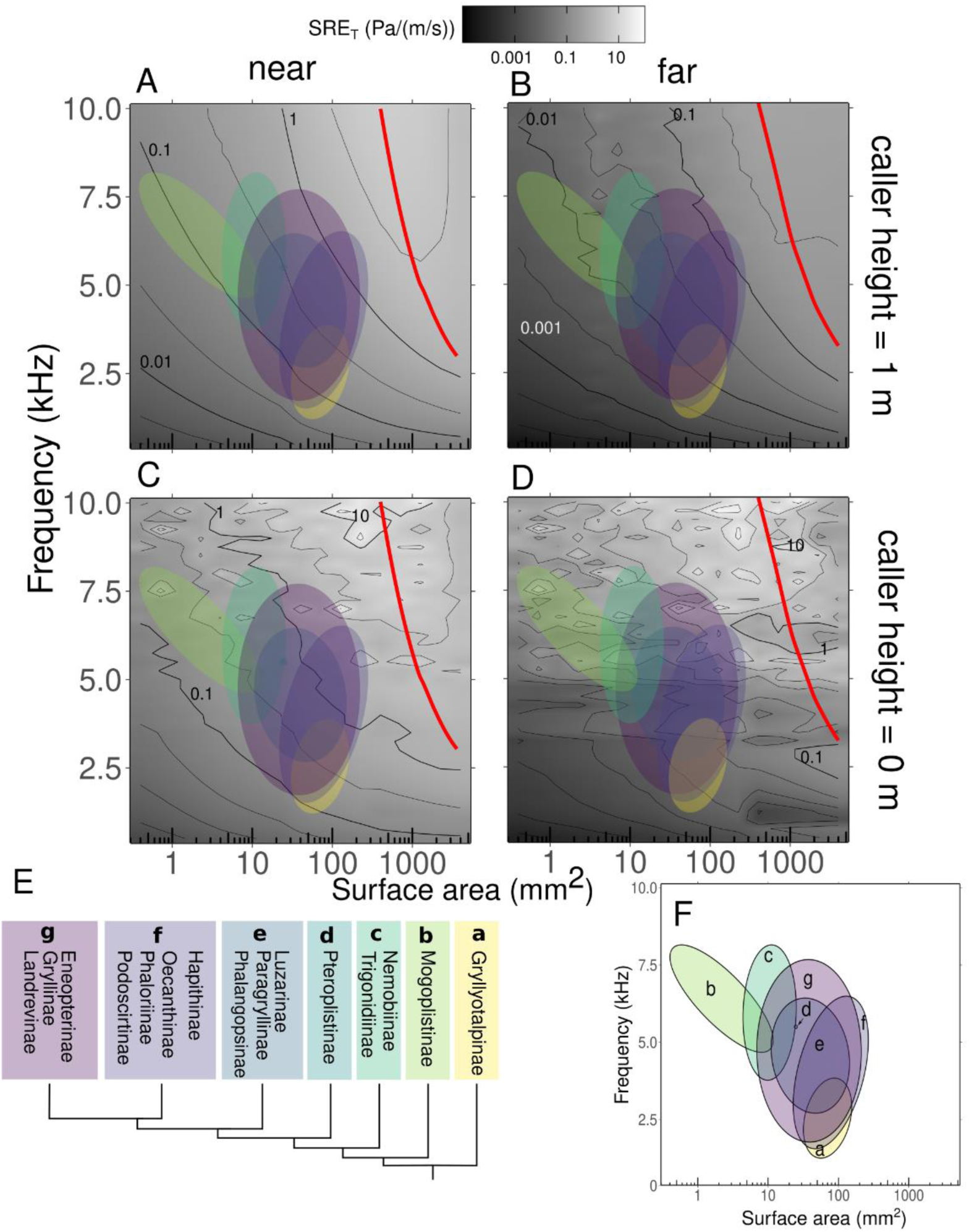
Sound propagation efficiency when the cricket sings near the bare hard ground is orders of magnitude higher than in free space. Each panel represents a combination of caller height above ground (0 m or 1 m) and receiver distance from caller (0.05 – 02 m “near” and 0.8 – 0.9 m “far”). **A.** Distance = near, height = 1m, **B.** Distance = far, height = 1 m; **C.** Distance = near, height = 1 m, **D.** Distance = far, height = 0 m. Red lines indicate the optimal radiator size, or the size at each frequency that would produce an ideally baffled calling scenario (see description of baffle calculations for SRE_V_). Note that the frequency range is reduced compared to Figure 1, in order to exclude high frequency callers which likely use alternative vibrational modes. Each clade of animals is represented by a colored ellipse. **E.** Phylogeny representing each clade **F**. Key to clade represented by each ellipse.

Next we plotted an SRE_V_ landscape (Fig 1) for the full acoustic-morphospace. On this landscape, we plotted the locations of the 112 species of crickets with known wing size and frequency allowing us to infer their SRE_V_ (Fig 1). These data therefore demonstrate where crickets lie in the efficiency landscape (Fig 1F), and how much they could gain through the behavioral means of using of an optimal baffle (Fig 1G).

Previous work examining four species of crickets and a small number of other insects determined that they each perform less efficiently than the theoretical maximum for dipole sound sources (9). In our larger dataset, there is a clear ridge of high efficiency running through the SRE_V_ landscape (red line in Fig 1C). This line indicates the radiator size at which short-circuiting is minimized and efficiency maximized at each frequency when the radiator is suspended in free space. Despite sampling species with a wide range of wing areas (0.4 – 258 mm^2^) and call frequencies (1.6 – 27.9 kHz), all species lie below this line. The efficiency distribution among crickets is somewhat bimodal (Fig 1F, G). The nine species with an SRE_V_ above 1 Pa/(m/s) all had calling frequencies above 14 kHz and belonged to the subfamily Eneopterinae, in clade g. Given the difference in their song radiation mechanics (35) we excluded these individuals from subsequent analyses (see methods). After excluding these high frequency callers, we found that SRE_V_ ranges from 0.02 to 0.67 Pa/(m/s), mean: 0.19 ± 0.01 SE, n = 103.

Next, we calculated how much efficiency each species could gain by using a baffle. A baffle decreases acoustic short-circuiting and is functionally similar to increasing radiator size (8, 12). The efficiency gained from an optimal baffle is the same as from an optimally sized radiator (8). Therefore, we use the line representing optimal radiator size to calculate how much each species could gain in efficiency by using an optimal baffle. Each species had a calculated “baffled efficiency” by noting where its call frequency intersects with the line. If animals were to continue using the same call frequency, but used an ideal baffle, each species stood to gain between 1.8 – 36 times (5 – 30 dB) above their baseline SRE_V_ (mean: 7.9 ± 0.41 times, 16 ± 0.48 dB, n = 103). Among those who stood to gain the most included animals in clade b, specifically in the subfamily Mogoplistinae (scaly crickets). These animals have very small wings (mostly under 5 mm^2^), suggesting a poor match between sound wavelength and wing size. On the other hand, the animal closest to line was *Madasumma affinis*, belonging to the subfamily Podoscirtinae in clade f. This animal has the largest wing at 258 mm^2^, however, even this animal stood to gain 0.5 Pa/(m/s) (5 dB or 1.6x increase) with the use of an ideal baffle. Taken together, these data suggest that all crickets could increase SRE_V_, and therefore, stand to benefit from use of a baffle.

### Ground calling emerges as an alternative strategy to baffle use in complex environments

While analysis of SRE_V_ suggests that all crickets should use baffles, this prediction is based on sound fields travelling in free space and over short distances. It is possible that efficiency advantages from baffle use become negligible as sounds interact with objects in the cricket’s local environment such as the ground or the vegetation. Many baffle users are low frequency callers, and it is also possible that higher frequency crickets lose all advantage from baffling in complex acoustic conditions. Either of these scenarios would lend support to the lack of utility hypothesis.

To address whether and how the efficiency landscape is changed by realistic calling conditions, we used boundary element modeling. Specifically, we used this method to add a “ground” component to our existing models, where the ground could have different characteristics including vegetation cover. In these models, sound can be reflected and dissipated by the ground and the effect of the vegetation is captured by an excess attenuation term based on empirical data (see supplementary methods). We used empirical measurements of ground impedance and although our modeled ground is flat and smooth, these measures should take realistic ground variability into account. We also varied the height of the caller above the ground (ground calling: 0 m, elevated calling: 1 m). We measured efficiency again by normalizing sound levels against radiator vibration levels (see methods). Sound levels were measured at two distances from the caller: near (averaged from 0.05 – 0.2 m away), and far (averaged from 0.8 – 0.9 m away). To simplify analysis, we always measured efficiency at the same height as the caller. To differentiate this measure of efficiency from SRE_V_, we call it sound radiation efficiency along a transect, or SRE_T_ (Pa/(m/s)).

The most striking and unexpected result from our analysis was that calling from the ground (Fig 2C, D) yielded much higher SRE_T_ than calling from one meter above it (Fig 2A, B). This is reflected in the efficiency landscapes by an average increase of about 5x (14.5 dB) across the entire acoustic-morphospace that we measured. Indeed, the highest SRE_T_ observed with a ground caller was 4 Pa/(m/s) (Fig 2C), two orders of magnitude higher than peak SRE_T_ with an elevated caller (0.06 Pa/(m/s), Fig 2A). This increase in efficiency is likely due to two phenomena. The first is partial baffling offered by the ground, which will prevent some acoustic short-circuiting. A second phenomenon is likely the ground effect, in which the pressure field that would normally propagate below the radiator is instead reflected upward from the ground and mixes with the directly propagated field. Here, we see that the ground effect leads to constructive interference between the direct, and the reflected and the ground wave and an increase in sound pressure levels (36). We find that even optimally baffled animals could gain an average of 2.8x efficiency (9.2 dB) along the measurement transect, by calling from the ground compared to elevated calling.

On the other hand, calling from far above the ground yields SRE_T_ values that are similar in level to SRE_V_ values calculated in the ideal free-field scenario modeled previously. At further distances, the values decrease as predicted by spreading in open air. Taken together, our models posit that ground calling and elevated baffled calling are two potential alternative strategies to maximize efficiency.

### Calling from the ground is still efficient when ground properties and vegetation are varied

All grounds are not equivalent and the increase in the net increase SRE_T_ may depend on their properties. For instance, soft grounds or those covered with vegetation would be much more dissipative and may eliminate the advantage accrued from ground calling. To test this possibility, we investigated whether this alternative strategy framework holds up when these properties of the environment are varied. We found few differences in SRE_T_ with different types of grounds (Fig S7). SRE_T_ tends to be slightly higher with a “soft” ground, which is better at dissipating sound, similar to a freshly tilled agricultural field (see methods) and this effect is magnified further away from the caller. This suggests dissipation has a higher effect further from the source, and primarily on propagation, whereas here the phase shift is more appropriate for constructive interference near the source. With a harder, more reflective ground, similar to a tightly packed forest floor, SRE_T_ is slightly lower. However, significant differences between these two ground types occur at wing sizes well outside the natural range for crickets. At close distances, and particularly above the ground, the differences between ground types are very small (Fig S7). Therefore, all future analyses assume a “hard” ground.

Finally, we tested whether vegetation would reduce the predicted SRE_T_ landscapes for ground calling. Vegetation does slightly decrease the magnitude of SRE_T_ overall and unsurprisingly, there is a slight frequency dependence where SRE_T_ is lowered slightly more at high frequencies (Fig S8). This suggests that high frequency callers may be at an increased disadvantage when calling in vegetation as suggested before (29), and will see diminishing returns when using a baffle. However, we found that excess attenuation due to vegetation does not significantly change the overall patterns of efficiency. By and large, it shifts the efficiency landscape to a lower point at most points within the parameter space (Fig S8) (37). However, the efficiency near the ground remains higher than the efficiency 1m above the ground (Fig S9). Finally, the effects of vegetation on SRE_T_ are undoubtedly more complicated than an excess attenuation factor. Modeling plants explicitly, at a variety of shapes and sizes, would be a useful extension to this study. However, since the efficiency of ground calling remains higher than calling from 1m above ground, we conclude that calling from the ground remains an effective alternative strategy, even if the ground is soft, or covered with some vegetation.

### Ground calling does not significantly degrade call directionality

So far, our analysis has used the loudness of calls to define efficiency. However, a call must be both loud and directional to be effective. That is, the call must present a spatial gradient that a potential mate can follow to the source. Previous data has suggested that such gradients are severely degraded in ground calling crickets (29, 33, 38, 39), but not in elevated calling (40). This suggests that SRE_T_ gains from ground calling may trade off against call directionality. To test this, we analyzed call directionality by designing a directionality metric that captured how difficult it would be for a female cricket to follow an acoustic gradient back to the call’s source (see methods). A value of one indicates that the gradient along a transect perpendicular to the wing planes is always in the “correct” direction, that is, sound pressure level increases as the female moves toward the caller in steps of ∼ 2 body lengths (2 cm). A lower value means that over some stretches of this transect, SPL increases and at other steps it decreases. A value of 0.5, for instance, means that the SPL decreases over 50% of the steps as the female moves closer.

We find that directionality varies with respect to frequency, radiator size, and height from ground (Figs S10, S11). Although ground calling does experience a loss of directionality compared to elevated calling, these losses are small. Near a ground caller, calls are all strongly directional (> 0.9) below about 5 kHz, except for very small wings. Further from the caller, calls are strongly directional below about 3.5 kHz. Therefore, high frequency callers would be most susceptible to the gradient effects. However, even below these cutoffs, directionality rarely drops below 0.5 in any condition, and ground calling remains a viable strategy. It should be noted that other studies have found more substantial degradations in call directionality in sounds traveling along the ground, but over greater distances than our current models (33). However, data for both field crickets and tree crickets suggest that the SPL of typical cricket calls drop below threshold at about 1 m from the caller (38, 40), and therefore we considered this a biologically relevant distance over which to measure directionality.

### Alternative calling strategies are likely in use by some cricket species

Based on the overall efficiency landscape, ground calling and baffled calling are potential alternative strategies to maximize efficiency. However, we have so far considered the full acoustic-morphospace, i.e. all possible combinations of radiator (wing) size and call frequency, but most of these combinations are not used by real crickets.

To shift our focus to the sampled cricket species, we tested where alternative calling strategies may offer the largest advantage to actual crickets. For each of the 103 animals in the dataset, we calculated SRE_T_ for each of three alternative strategies as measured far from the caller: calling from the open hard ground (ground calling), and from within vegetation 1 m off the ground with no baffle (unaided calling), and from within vegetation with a baffle (baffled calling) (Fig. 3a). We compared both baffled calling and ground calling to unaided calling as a baseline. It would be ideal to determine whether animals, in fact, use the strategy that we predict should maximize efficiency based on known calling preferences. Unfortunately, we do not have data on calling preferences of many sampled animals. Below, we give four examples of species where information about calling behavior is available.

**Figure 3.**
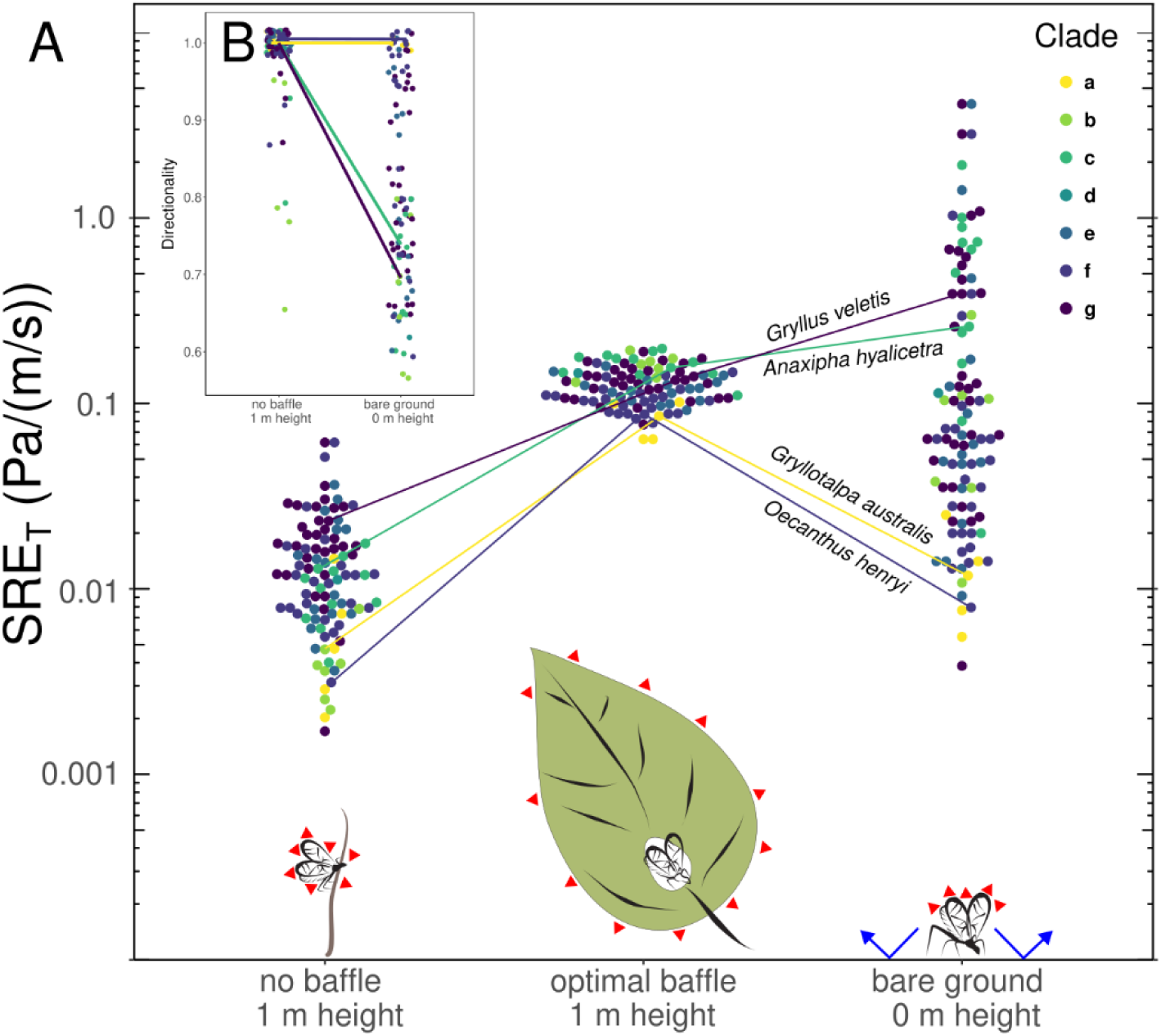
The calling strategy with highest efficiency varies depending on species, whereas directionality does not decrease greatly even with ground calling. Four representative species are illustrated with lines. **A**. Comparison of SRE_T_ on bare ground with no baffle, and 1 m in vegetation with and without an optimal baffle. These measurements were taken far from the caller, i.e., an average of the SPL at a distance of 0.8-0.9 m from wings. Images below are not to scale and show a schematic representation of the three different calling strategies. Red arrows represent the location of acoustic short-circuiting. Blue arrows show sound reflection from the ground. In the optimally baffled scenario, the baffle effectively increases the size of the radiators. In the ground calling scenario, the ground provides some reduction of acoustic short-circuiting and halves the air volume within which sound is radiated. It simultaneously reflects sound upwards, allowing recapture of a significant amount of acoustic energy through constructive interference. Both strategies improve efficiency to different degrees depending on the combination of wing size and call frequency. **B.** Optimally baffled and ground calling do not substantially differ with respect to directionality. Points are jittered slightly for clarity. The optimally baffled condition all had a directionality of 1, so this condition is not shown. Note that the y-axis begins at 0.5.

We start with *Oecanthus henryi*, a well-studied species of tree cricket in clade f. This species stands to gain efficiency on the order of about 2.5x, (8 dB) from ground calling compared to unaided calling according to our data (Fig 3A). However, they could gain 28x, (28 dB) if they baffled. Indeed, what is known of this species’ natural history bears out our predictions; *O. henryi* not only calls from vegetation, including vegetation that is suitable for baffle building and use (6, 40), in fact, this species is well known for its baffle-building behavior (7, 10, 12). For *Gryllus veletis*, a field cricket in clade g, on the other hand, we predict the opposite. On average, ground calling gives an advantage of 16x (24 dB) above unaided for this species, whereas baffling gives an advantage of about 5x (15 dB) above unaided. Again, behavioral data suggests that many field crickets (including this species) indeed prefer to call from ground habitats that we predict would maximize their efficiency (41). *Anaxipha hyalicetra*, a sword-tailed cricket from clade c also stands to gain more from ground calling (20x, 26 dB) than from using a baffle (12.5x, 12 dB). This species is known to call from the surfaces of fallen leaves and bunch grasses (42), suggesting that other objects in the environment may act as a “ground”.

Finally, *Gryllotalpa aulstralis*, a species of mole cricket in clade a, represents an interesting exception to this alternative strategy framework. This species stands to gain a great deal from baffling (18x, 25 dB), compared to ground calling (2.5x, 8 dB). Yet, this species (and all species in clade a) are known to exclusively call from the ground and do not use baffles. However, they do use an acoustic aide. Mole crickets build and call from burrows which function as resonators and convert them into monopole sound sources, eliminating acoustic short-circuiting through a different mechanism than baffling (28, 43). Indeed, it is possible that other acoustic means of maximizing call efficiency exist and could in the future add further complexity to our hypotheses.

Finally, we find if animals baffled, but their calls propagated though vegetation compared with no vegetation, the gains would be relatively small in most cases (< 6 dB SPL) (Fig S9), We also performed a similar analysis for call directionality (Fig 3B, Fig S11). However, since directionality was quite high for all calling conditions, we suggest that directionality does not preclude one alternative strategy over another.

## Discussion

### Why would baffle use evolve among crickets in the first place?

From our data, exploiting the ground effect by calling from the ground emerges as a viable alternative to tool use in crickets. This simple site selection strategy can even exceed the efficiency gains of tool use in some scenarios. Given that making and using tools like baffles requires a specialized behavioral repertoire, and precise execution of these behaviors (7, 12), the real question becomes why a species would ever use this strategy if simpler site selection preference for the ground could give similar increases in efficiency.

There is evidence that crickets have been singing as early as the Cretaceous period (44). These early calling crickets were likely ground dwellers, with some species subsequently moving up into vegetation as the group diversified (45). We therefore suggest that baffle-using crickets may have originally moved up into vegetation for non-acoustic reasons, whether it was to exploit additional food resources or avoid predators. Baffle use would have then evolved secondarily to recover some of the efficiency lost when abandoning ground calling. The biophysical modeling methods presented here open the door to testing such a hypothesis about baffle use.

It is also known that crickets call from other “ground” substrates such as tree trunks, cave walls, or artificial structures (25, 28), which our data suggest could also increase calling efficiency (35, Fig 4C, D). The efficiency effects of these substrates could be further investigated using biophysical models contributing to our knowledge of acoustic ecology. In principle, we could even model the wings of extinct crickets, and estimate calling frequency based on the stridulatory apparatus on the wing (46, 47). By bringing extinct crickets “back to life” in this way we could ask questions about the evolution of acoustic tool use and calling ecology more broadly. We suggest that biophysical modeling, grounded with data from real animals, can be a valuable tool for any biologist wishing to better characterize and understand tool use in the context of animal communication.

**Figure 4.**
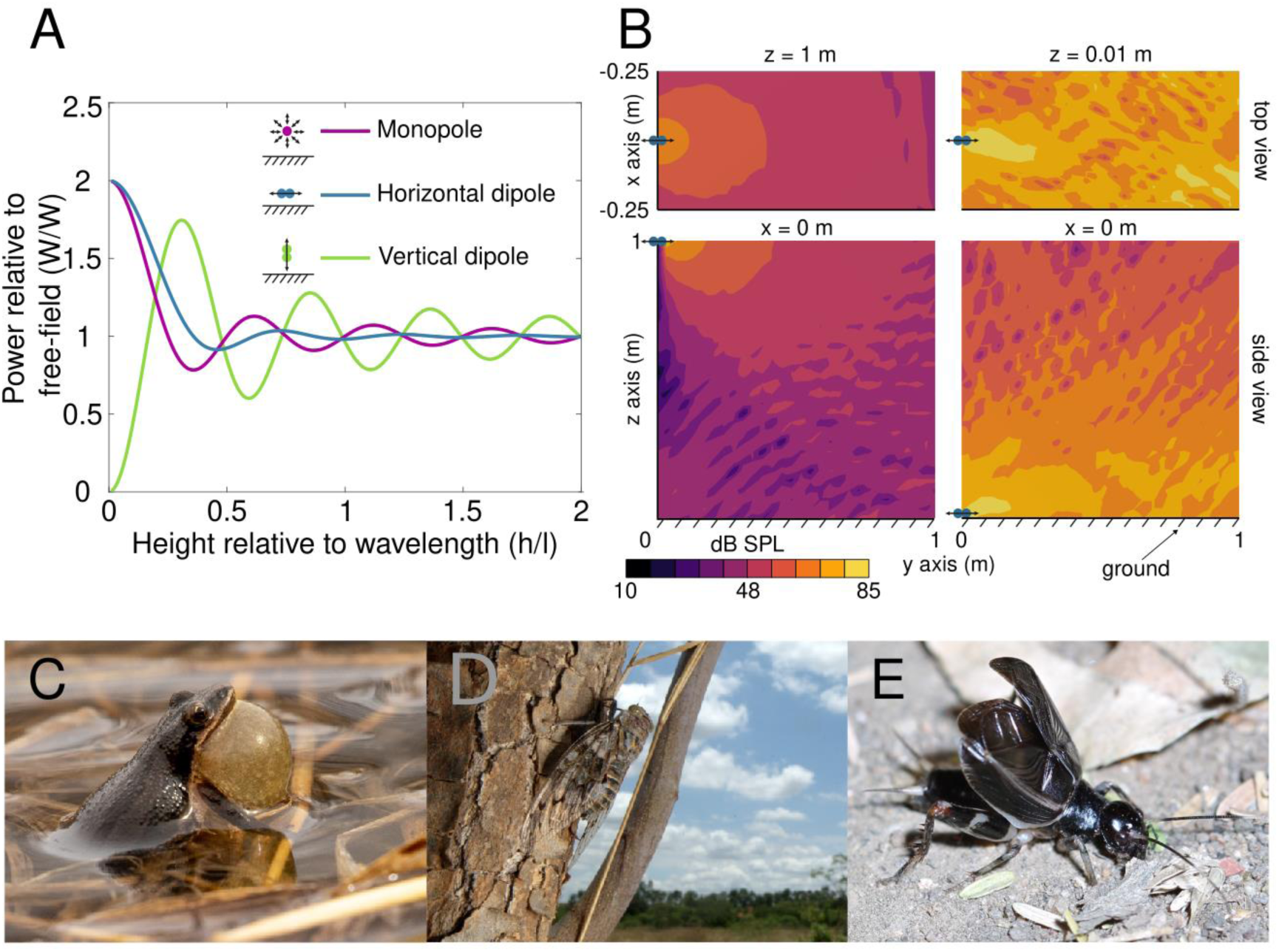
Hard reflective surfaces like the ground or water offer all acoustically active animals an increase in sound radiation efficiency. **A.** The presence of a perfectly reflective surface near an acoustic source can lead to an increase of the power radiated in the hemisphere above the source, relative to the power that same source would radiate over a full spherical volume in free space. Both monopole and horizontal dipole sources (vibration axis parallel to the reflective surface) can get a large boost in power when they are close to the reflective surface, relative to the wavelength of sound being radiated. (Reproduced from equations in ref. (54)). In contrast, vertically oriented dipoles need to be further from the reflective surface to get an equivalent gain in efficiency. **B.** In fact, near the ground, the sound field from a dipole source is expected to become less directional and we observe this in our models. Data shown here are from a radiator of surface area of 126 mm^2^ and at frequency 5 kHz, which is seen to be directional 1 m above the ground, but much less so when right next to the ground. **C-E.** Different animals, especially small animals who have low sound radiation efficiency whether they are monopole or dipole sources may call from near acoustically hard and reflective surfaces, like **C.** water, **D.** tree trunks, or even the **E.** ground may be exploiting this effect. (Photos from left: Doug Mills, Natasha Mhatre & Brandon Woo). Animals that are monopole-like sources are likely to orient themselves parallel to the surface since this is advantageous for camouflage as well. However, crickets (dipole-like acoustic sources), are constrained to expose themselves by calling with their wings perpendicular to the ground to maintain the advantageous horizontal dipole orientation.

### Sound radiation efficiency in animal acoustic communication

In the field of animal acoustic communication, calling from the near the ground has been thought of as severely disadvantageous, whether in insects, birds or even primates (27, 28, 48–53). One reason for this is that reflections from the ground degrade temporal cues in calls and songs (31). However, most animal calls are repetitive and redundant, since they are used for functions such as sexual or territorial signaling, or as contact or alarm calls. Here, fine temporal structure is not crucial and the primary functional factor is how loud the sound is (28). How then is loudness affected by the presence of a reflective surface such as the ground? In animal acoustics, it is widely believed that the ground generates high amounts of excess attenuation compared to simple spherical spreading of sound (27, 28, 48–53). Yet, many animals call from the ground, tree trunks, or other vertical surfaces such as cave walls in many ecological contexts (28).

Previous acoustic analyses have typically measured excess attenuation as observed at two distances from the source. Thus, these analyses only account for propagation losses and fail to consider the first step in the process, sound radiation efficiency, i.e., the relationship between the vibration amplitude of the radiator and the level of the sound field that is generated. Here we show that for dipole singers like crickets, calling from the ground can actually provide a substantial boost in sound radiation efficiency, outweighing propagation losses at distances and receiver positions relevant to cricket behavior. It is only when we take this first step in the process of sound generation into account, that it becomes clear that calling from the ground may be an advantage rather than an impediment.

In fact, this phenomenon may be much more general than currently appreciated; analytical findings from as far back as 1957 show that the acoustic power radiated by monopoles and horizontal dipoles (such as cricket wings) increases as they get closer to a perfectly reflecting surface, by as much as a factor of two (Fig 4) (54). When acoustic sources are close to the ground, the reflected sound field effectively sums with the direct sound field, and at very close distances, this summation is near perfect. In particular, the radiated field from dipoles becomes almost monopole like when the source is close to a reflective ground (Fig 4) (55).

Of course, real grounds are not perfectly reflective and do have a finite impedance; they absorb some sound energy (36, 56). However, this does not significantly alter the theoretical expectation of increased sound radiation, except that an additional ground wave is formed in addition to the reflected wave (36). Nonetheless, dipole and monopole sources near the ground are still theoretically expected to have higher sound radiation efficiencies compared to the free or direct field condition (36, 57). In our analyses, we considered dipole sound radiators of finite size, above realistically parametrized grounds. We found sound radiation efficiency increases considerably near the ground. In fact, in some cases efficiency increases by even more than a factor of 2, likely through the baffling effect offered by the ground against acoustic short circuiting.

These findings extend beyond crickets, as they are true for both horizontal dipoles and monopoles (Fig. 4). All acoustically communicating animals are considered to be either dipole or monopole-like sound sources. Among non-cricket invertebrates, both fruit flies and mosquitoes produce sounds with their wings, functioning as dipoles (9). Other invertebrates that use tymbal-based sound production like cicadas and wax moths behave like monopoles (9). Among vertebrates, the sound fields produced by bats are well captured by a baffled-dipole piston model (58, 59) and whales have a similarly directional sound field (60). Most acoustically active vertebrates such as fish, frogs, reptiles, birds and mammals are considered to be monopole sound sources (27, 61).

Thus all these animals can exploit this mechanism for gaining efficiency (9, 27, 61). In effect, by shifting the focus from sound propagation to sound radiation efficiency, we posit an explanation for both the rarity of tool use and high number of animals that call from the ground. Indeed, our findings, and a reconsideration of established acoustics theory, leads us to the exciting discovery of a hitherto unknown mechanism for increasing calling efficiency available to all acoustically communicating animals.

## Materials and Methods

### Specimen Data

We collected data on wing surface area and call frequency for 112 cricket species distributed across 7 clades *sensu* Chintauan-Marquier et al. (34) (Fig 1, Fig S1). We restrict our analysis to Grylloidae, since they are known to raise their wings when singing (62). This behavior means that they are dipole sound sources, and acutely affected by acoustic short-circuiting (5, 9). Data were obtained from a variety of databases including Orthoptera Species File (56), Crickets North of Mexico, and numerous publications (all references are available in Tables S1, S2). For a few species of Oecanthines, wings were provided by Nancy Collins and photographed in the lab under a dissecting microscope. We measured surface area of the entire left forewing including the lateral field. Fitting an ellipse to the wing, we calculated aspect ratio (length of ellipse/width of ellipse). All image data were gathered using ImageJ version 1.53 (64). The fundamental frequency of cricket advertisement calls was captured using Raven Lite version 2.0 (Cornell Lab of Ornithology, 2020). In crickets, call frequency typically does not change with environmental conditions (66), in a few exceptional groups such as tree crickets this change is typically <1kHz (67) which we do not expect to change our conclusions strongly. Therefore, we do not consider recording conditions and treat song frequency as an invariant character of cricket species. For our models, we assume that call frequency is identical to wing oscillation frequency as is widely accepted (68). Both have been simultaneously measured in at least two species of crickets where this has been verified (69, 70).

When multiple specimens of a single species were analyzed, averages were used for wing size and call variables. To better represent the full range of wing size and call frequency in our dataset, we included some specimens in the histograms showing wing size and frequency (Fig 1D, E) for which we had only one type of data. Twelve animals had only wing size, but not call data, and 57 animals had call, but not wing size data (see Tables S1, S2 for details).

### Finite Element Models for Sound Radiation Efficiency

We estimated the sound radiation efficiency of crickets calling in open air using finite element (FE) analysis (Fig S2). All models were built in COMSOL Multiphysics version 5.5. All models used the pressure acoustics module and were solved in the frequency domain assuming a steady state. The Helmholtz equation was the governing equation.

#### Model geometry, boundary conditions, symmetry, and vibration

Animals were represented by two ellipses which modelled the forewings sitting next to each other along the long axis, in the same plane (see Fig S2 for description of symmetry). The size and shape of the radiator and call frequency determines radiation efficiency, and not the radiator’s material properties (8). Surrounding the wings was a 20 cm radius spherical acoustic domain consisting of air. We applied a time- and space-averaged velocity normal to the entire wing surface at 0.13 m/s. This was the value measured from the wings of singing *Oecanthus henryi,* the only known estimate for crickets (7). However, for our analysis we use efficiency rather than sound pressure levels. Efficiency normalization allows comparison between species. We vibrated the wings from 0.5 - 32 kHz, in increments of 0.25 kHz. SRE_V_ was calculated from model outputs as a volumetric average of the absolute pressure in the acoustic domain, divided by the time- and space-averaged velocity of 0.13 m/s, resulting in units of Pa/(m/s).

#### Finite Elements

3D tetrahedral elements were used in both the acoustic domain and PML. After undertaking a mesh size sensitivity study (Fig S5), we chose the “extra fine” mesh setting in COMSOL, with about 60000 elements in the acoustic domain. This number did vary somewhat with wing size, as fewer elements are used with very small wings.

#### Model Parameters

We ran the FE model at wing surface areas from 0.4 – 4000 mm^2^, scaled logarithmically by the equation 4 × 10^x^, where x ranges from −1 to 3). We used an aspect ratio of 2 (wings are twice as long as they were wide). Our chosen aspect ratio of 2 was within the range of most cricket species (median: 1.7, range: 0.7 - 3.7). Aspect ratio did not greatly alter efficiency, except at aspect ratios > 5 (length of wing 5x the width), which were not observed in real wings (Fig S4). For natural aspect ratios, differences in SRE_V_ at a given wing area and frequency never exceeded 3 dB.

#### Other Modeling Considerations

The cricket body was not included in our models as it was found to have negligible effects on SRE_V_ at all wing sizes and frequencies (mean difference: 0.05 ± 0.01 dB). We also evaluated whether applying vibration to only a part of the wing (a “harp”) influenced sound production. Some cricket species (though not all) use this sound production method (71). We found only minor increases in SRE_V_ between vibrating only a harp or vibrating the entire wing (mean: 4 ± 0.08 dB), except at wing sizes well outside the range of the real wings (Fig S4).

### Boundary Element Models for SRE_T_

To test hypotheses about how cricket calls interact with objects in the environment, we needed to include an additional domain in the model: a “ground” with realistic parameterized acoustic impedance. To make this model as realistic as possible and to minimize boundary effects, we needed to make the ground element as large as possible relative to the size of the wings. The combination of the large size of ground and the high frequencies of interest resulted in finite element models that were too computationally intensive to run. We therefore turned to boundary element (BE) modeling as an alternative means of assessing SRE_T_.

Boundary element models and finite element models are both numerical methods for solving the Helmholtz equation to capture a developing sound field within a medium. However, they discretize space within the model differently. FE models discretize volumes by partitioning into a 3-dimensional mesh of finite elements. BE models on the other hand reduce computational cost by discretizing only the boundaries of the acoustic domain and assume a linear homogenous medium in all other spaces. The BE formulation trades off some specificity in exchange for computational efficiency, allowing us to make large, biologically relevant models to assess sound SRE in a spatially explicit manner.

We ran our BE models using the pressure acoustics, BE module in COMSOL. All models were run in the frequency domain and assumed steady-state behavior. The Helmholtz equation does not take attenuation due to damping into account, which can become an issue at distances far from the source. However, at frequencies >500 Hz, attenuation due to damping is only about 2 dB per kilometer (32), so we considered this effect to be negligible over the distances of interest for this study (0.2 – 1m).

#### Model geometry, forcing, and boundary conditions

Wings in the BE model were modeled in the same way as in the FE model (Fig S2), with no material properties and one-way coupling between the wings and sound fields. The ground was modeled as a rectangular slab, 0.5 m wide, 2 m long, and 0.10 m thick. Wings were positioned perpendicular to the top surface of the ground, with the flat surfaces of the wings aligned with the short axis of the ground. The wings were centered with respect to ground. The wings were placed above the ground at either 0 m, or 1 m. The same time- and space-averaged velocity was applied, and the same set of wing surface areas were used. A sound-hard boundary was applied to the bottom surface of the ground slab. Because we were interested in spatially-explicit measures of efficiency as sound propagates across ground, we did not use symmetry conditions to create this model. However, because the sound fields should be symmetric on either side of the wings, we only measured the sound field on one side.

#### Model Parameters

We used a restricted frequency range for the boundary element models, ranging from of 0.5 – 10 kHz, in increments of 0.25 kHz. We chose 10 kHz as the cutoff because very few animals call above this frequency, and those that do were Eneopterine species who were likely using a vibrational mode inconsistent with the piston mode that we have implemented (35). High frequency (> 10 kHz) callers were included in the finite element models to give a general sense of where they might fit in with the other animals, but in reality no animals occupy this space in the landscape and all analyses explicitly comparing species exclude them.

To model how the ground interacts with sound, we applied an acoustic impedance to our modeled ground. Acoustic impedance quantitatively describes how much sound energy is dissipated by the ground, compared to the energy reflected. We used the Attenborough slit-pore model to implement ground impedance. This model uses three parameters to capture both dissipative and reflective properties: flow resistivity, pore density, and porous layer depth. We modeled two different types of ground, a “soft” ground (flow resistivity: 2000 kPa×s/m^2^, porosity: 0.6) which is less reflective and a “hard” ground (flow resistivity: 9 kPa×s/m^2^, porosity: 0.4), which is more reflective. Porous layer depth was held constant for both treatments, at 0.04 m. The two parameter sets were taken from empirical measurements of a “soft” freshly-tilled field and a “hard” forest floor (56).

#### Transect sound radiation efficiency

In the FE models, we calculated a volumetric average of absolute pressure within the acoustic domain. However, this measure would not be appropriate to assess sound radiation efficiency at a distance, as the sound waves’ interactions with the ground would accumulate as distance from the source increases. Therefore, we calculated SRE_T_ in a spatially-explicit manner. We measured absolute pressure at 50 points along a 1m long line parallel to the long axis of the ground, at the same height as the wings. The line originated at the center between the two elliptical ‘wings’. We divided this line into “near” and “far” from the caller: near = 0.05 – 0.2 m from wings, far = 0.8 – 0.9 m from wings. Efficiency was calculated by dividing sound pressure level (Pa) by 0.13 m/s, the space and time averaged velocity applied to the wings. We also created an additional boundary element model with no ground, to allow for direct comparisons between ground and no ground and to sanity check the BE method compared with the previous finite element models.

#### Finite Elements

Tetrahedral elements were used on the surface of the ground and 2D triangular elements on the wings. After performing a similar sensitivity study as with the finite element models, we decided on a maximum element size for the wing surfaces of 0.5 cm and 1 cm for the surface of the ground. Since the sound wave is not explicitly modelled, this element size is not related to sound frequency, and instead captures boundary conditions and hence can be larger than in the finite element models.

### Excess attenuation due to vegetation

To calculate the effect of vegetation, we used existing models to calculate how standing vegetation is expected to impact call efficiency. We then subtracted this excess attenuation from the COMSOL result. We calculated excess attenuation using the following empirically derived equation(37):

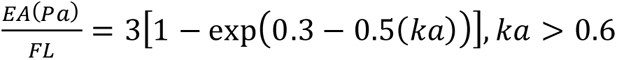

Where EA(Pa) represents excess attenuation due to foliage, F = foliage area per unit volume, L = path length, k = wavenumber, and a = average leaf size. We used values empirically derived for dense foliage with relatively large leaves (37), such as those used to construct baffles in known baffle-using species. To approximate the foliage area and leaf size that a typical baffle user would prefer, we used F = 6.3m^-1^ and a = 0.0784m in our measures of excess attenuation.

### Efficiency of individual species and how much they gain with baffle

To test the lack of utility hypothesis, we used the output of our models to estimate the gains in efficiency that each species could attain if it used an optimal baffle. The efficiency of a radiator in an optimal baffle is the same as the efficiency of an optimally sized radiator for which short-circuiting is minimized and efficiency is optimized in free space (8) (Fig S6).

We identified this size for both the idealized measure of efficiency (SRE_V_) and the more realistic scenario involving a ground and vegetation (SRE_T_). For each modeling scenario, we estimated the efficiency of each cricket species in our dataset, given their wing area and call frequency. Next, we calculated the quantity ka for each surface area-frequency combination in the model, where k is the wavenumber and a is the effective radius of the sound radiating plates (8). ka is a dimensionless quantity used in acoustics, which helps define when a radiator of a particular size transitions from being inefficient sound radiator at low frequencies to an efficient high frequency radiator (an optimally sized circular piston has ka = 1) (8). However, the radiators being considered here are two aligned ellipses and at their particular value of maximal efficiency ka will be different. To estimate optimal ka for cricket wings, we plotted ka versus efficiency as measured from our models, with a separate trace for each frequency (Fig S6). We then identified the ka at which maximal efficiency was reached for all frequencies. In our FE models, we found optimal ka to be about 1.3. For the boundary element models at the far distance, optimal ka was approximately 1.55 (Fig S6).

Next, we performed a simple linear regression between frequency and maximal efficiency at optimal ka, then calculated the slope and y-intercept of this regression (Fig S6). We used this equation to calculate optimal baffled SRE_V_ and SRE_T_ for each species. The relationship between frequency and efficiency differed depending on condition (open ground vs ground + vegetation) (Fig S6), so this regression was performed separately for each environmental condition when calculating optimal baffled efficiency for a given condition. It should be noted that due to modelling constraints we were only able to calculate baffled efficiency on the ground for animals with a carrier frequency >3.5 kHz.

### Directionality index

To address how difficult it would be for a female to localize a male call, we assessed the directionality of the call in each modeling scenario. In an open sound field with no ground, the sound level is expected to decrease smoothly following the inverse square law (72, 73). A cricket moving toward the source of a call should therefore always experience either an increase in loudness, or, if the increase is below the animal’s difference threshold, no change in loudness. A cricket should always move in the direction of increasing SPL to locate the singing male. However, in reality, sound fields become more complicated when they interact with the ground, resulting in a noisy relationship between SPL and distance (74). In such sound fields, female phonotaxis may fail as there is no clear acoustic gradient to follow to the source. To quantify this degree of potential “confusion”, we calculated a directionality index for each modeling scenario. First, we calculated Δ SPL between each two adjacent points 2 cm apart (∼ 2 body lengths for most animals in this analysis). Δ SPL was calculated starting at 1 m away and moving toward the source. Next, we classified each of these values as either consistent with expected change in SPL or inconsistent. Consistent values represented either an increase, no change, or a decrease smaller than Δ 3dB SPL (a factor of about 1.4), thought to be close to the detectable threshold for crickets (75). See (74) for a more complex treatment of such thresholds. For our purposes, inconsistent values represented a decrease in SPL greater than 3 dB. For each modeling scenario, we calculated the proportion of Δ SPLs classified as consistent. This resulting value we call “Directionality” ranging from 0 to 1 (Fig S10). We calculated directionality for two different distance treatments, “near” was calculated from 0.05 – 0.2 m from the wings, and “far” was calculated from 0.5 – 1m from the wings.

## Data availability

Data used to create the figures can be found at DOI:10.5061/dryad.v15dv420b

## Acknowledgments

We wish to acknowledge a number of undergraduate students who assisted with the collection and databasing of cricket acoustic and morphometric data: Morteza Al Rabya, Nancy Kim, Shanker (Matthew) Nadarajah, and Daniel Xie. Nancy Collins provided specimens for several of the wing measurements. Graduate students Hossein Asgari and Reese Gartly assisted with multiplexing dozens of model runs on multiple machines. Emine Celiker and Damian Elias provided feedback on an earlier version of this manuscript. This work would not have been possible without an extensive worldwide network of specimen databases. We especially wish to thank those who contribute to, curate, or maintain Orthoptera Species File, Singing Insects of North America, Museum D’Historie Naturelle, and numerous university insect collections. We also wish to thank the photographers who graciously provided their images free of charge for inclusion in this manuscript. Finally, we would like to thank three anonymous reviewers whose comments greatly improved this manuscript.

We would also like to thank the following funding sources for their support of this research: NSERC Discovery (Grant no. 687216), and early career supplement (675248), and an NSERC Canada research chair (Grant no. 693206) to NM; the Western SEED grant to NM; the Western Postdoctoral Fellowship to EEB; Western Undergraduate Work Study to SD and HW; Western Undergrad Summer Research Internship to HW.

