## Supplemental Appendix for "The ground offers acoustic efficiency gains for crickets and other calling animals"

Supplementary Material

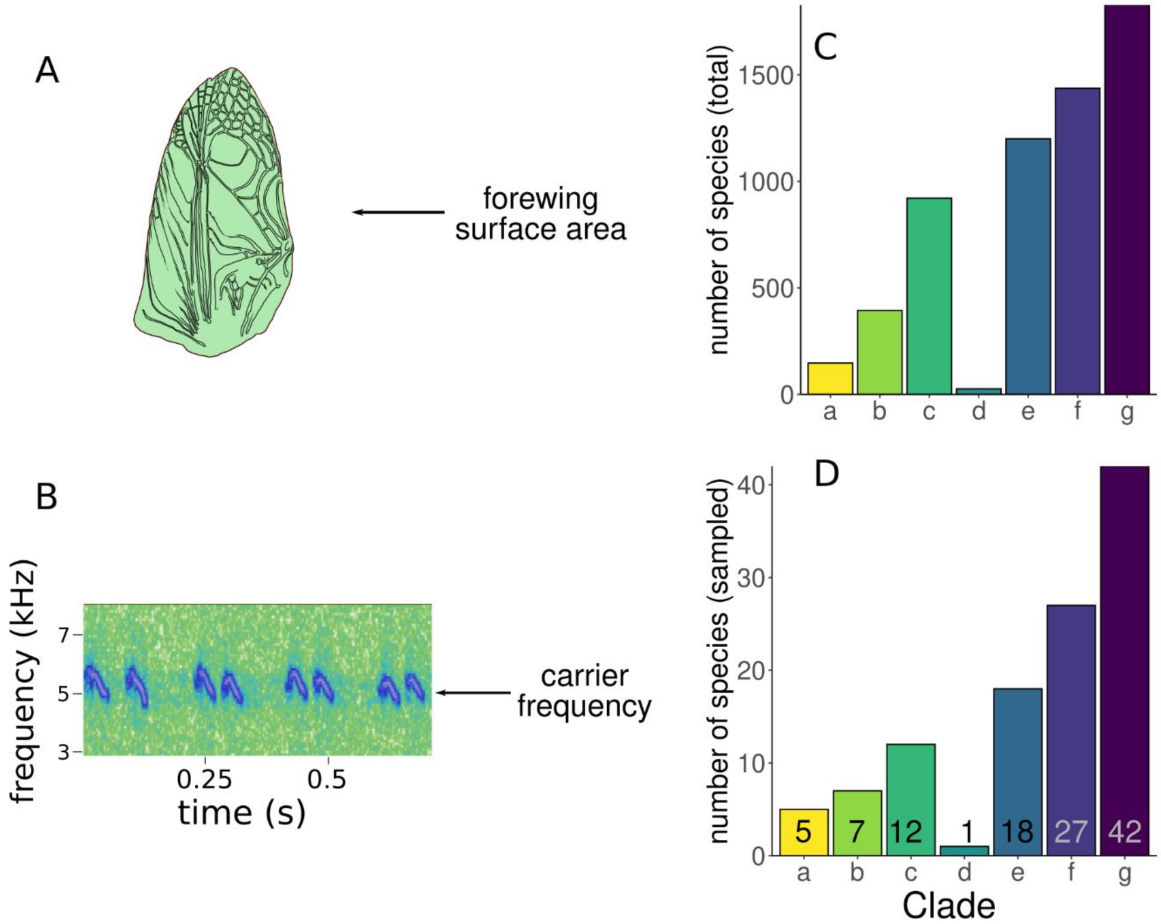


**Figure S1.** Measuring the acoustic-morphospace of crickets. Wing area and call frequency were quantified using published images and songs or spectra (see methods). **A.** Wing surface area was calculated as the area of the entire forewing. **B.** The carrier frequency (sometimes called fundamental frequency) of the call was identified from spectrograms. If the fundamental frequency occupied a sweep over a range of frequencies (as shown), an average was taken. **C.** Specimen sampling scheme. Distribution of all species in Orthoptera Species File (February 16, 2022) across each of the seven clades in Grylloidea. **D.** Distribution of species that were sampled for this study. Species numbers are given above clade labels.

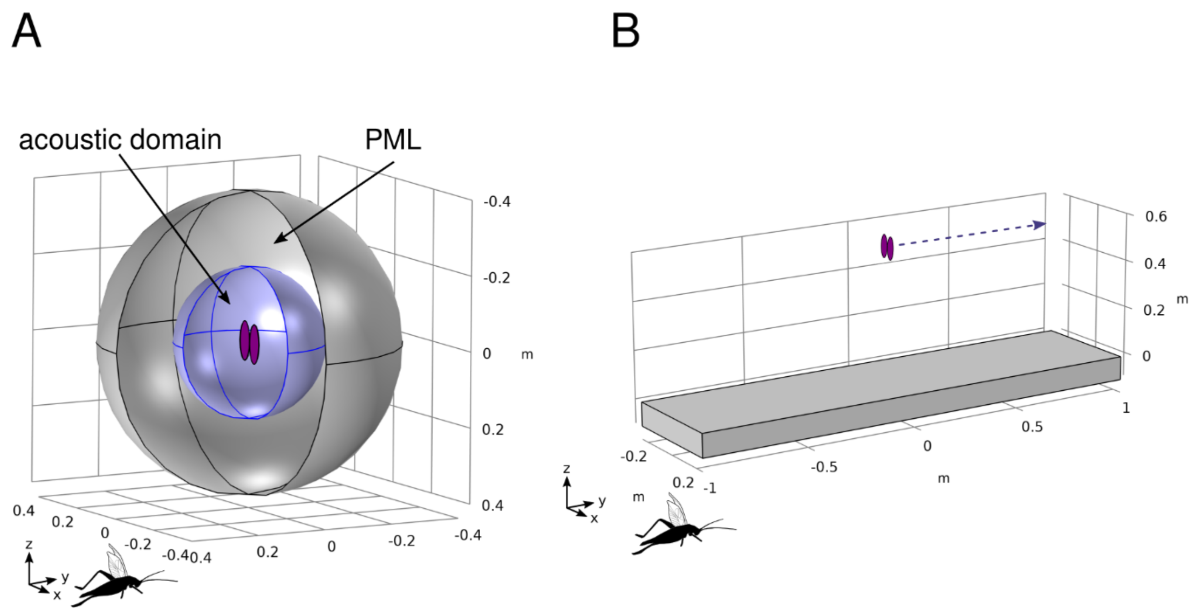


**Figure S2.** Geometry of biophysical models. **A.** Geometry of finite element (FE) models. The inner blue sphere is the acoustic domain. The wings are represented by purple ellipses in the center and the outer sphere is a perfectly-matched layer (PML), to mitigate boundary effects caused by the finite acoustic domain. Sound pressure level (SPL) is averaged over the inner sphere for the calculation of $SRE_V$. Wings vibrate in the piston mode along the y axis, as indicated by the cricket silhouette. **B.** Geometry of boundary element (BE) models. Grey shape represents a ground of defined acoustic impedance. Purple ellipses represent wings. Dotted line illustrates the line (transect) along which measurements were taken to assess $SRE_T$.

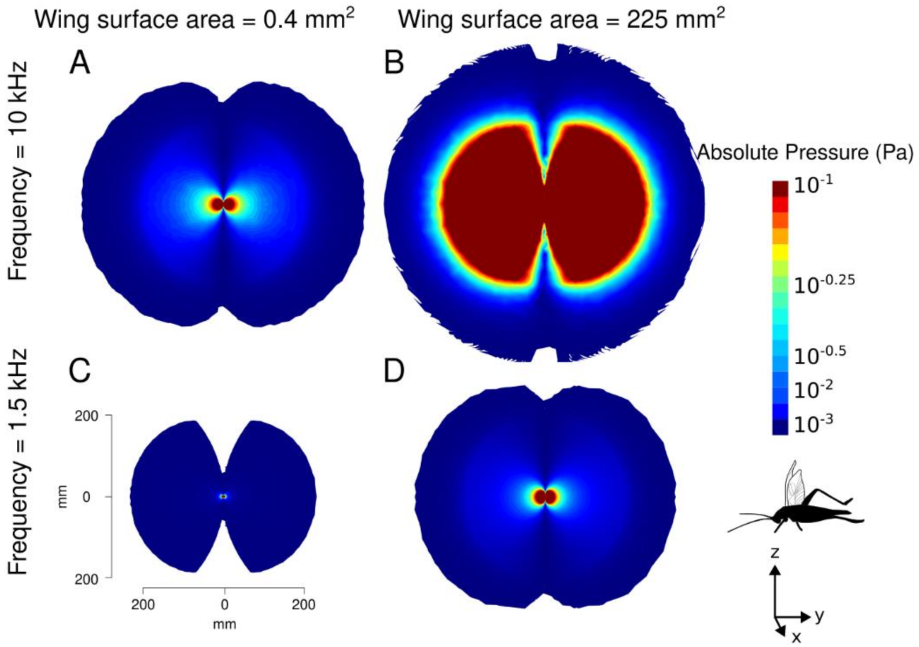


**Figure S3.** Sound fields produced by modeled wings vary depending on call frequency and wing size. Fields are oriented such that wings are perpendicular to page and vibrate left to right, as indicated by the silhouette cricket. Sound fields are given for the following combinations of wing size and frequency: **A.** wing size = 0.04 $mm^2$, frequency = 10 kHz; **B.** wing size = 225 $mm^2$, frequency = 10 kHz; **C.** wing size = 0.4 $mm^2$, frequency = 1.5 kHz; **D.** wing size = 225 $mm^2$, frequency = 1.5 kHz. Spatial scale given in C applies to all sound fields. SPL (here, indicated by the size and color of sound field) increases with improved match between wavelength of sound and size of radiator (wing) therefore so does efficiency. Cricket wings are usually small and this match is poor except at the extreme high end of radiator size and call frequency.

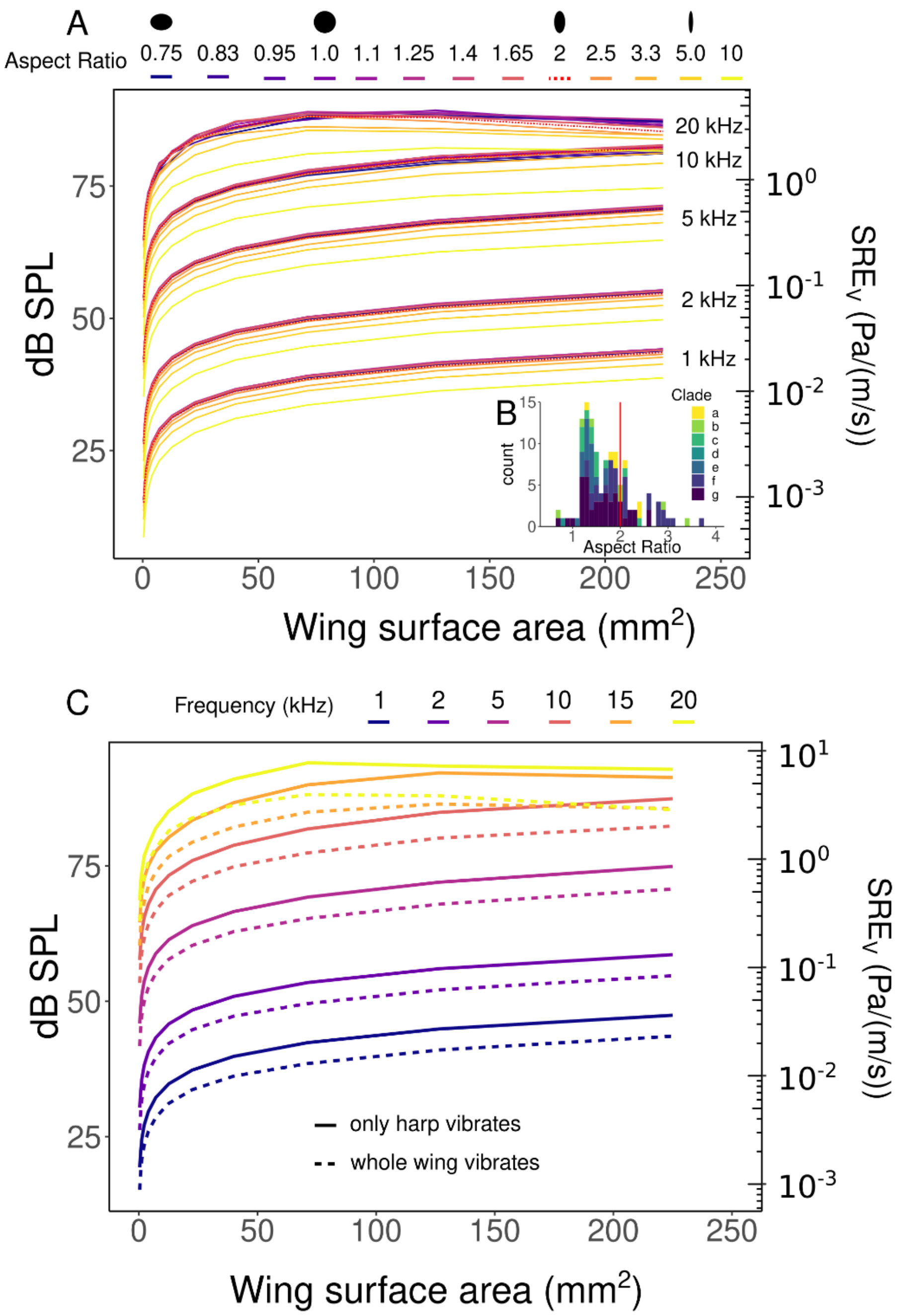


**Figure S4.** Wing aspect ratio and use of a “harp” resonator do not significantly impact $SRE_V$ within biologically relevant ranges of wing size and call frequency. **A.** The effect of wing aspect ratio on $SRE_V$ at six different call frequencies. The aspect ratio that was used on all models in this study (AR=2) is shown by red dotted line. **B.** Actual distribution of aspect ratios among species. Red line indicates aspect ratio that was used in our models (2). We see that while aspect ratio influences $SRE_V$, this effect is minor within the realistic range of aspect ratios (typically <3 dB for ARs from 1 to 3.3). **B.** The effect on $SRE_V$ of vibration spread over a small area (harp) compared with the whole wing. Some species of crickets restrict the vibrating portion of the wing to a “harp” region. However, we find that this does not strongly affect $SRE_V$ at any frequency within our range of interest.

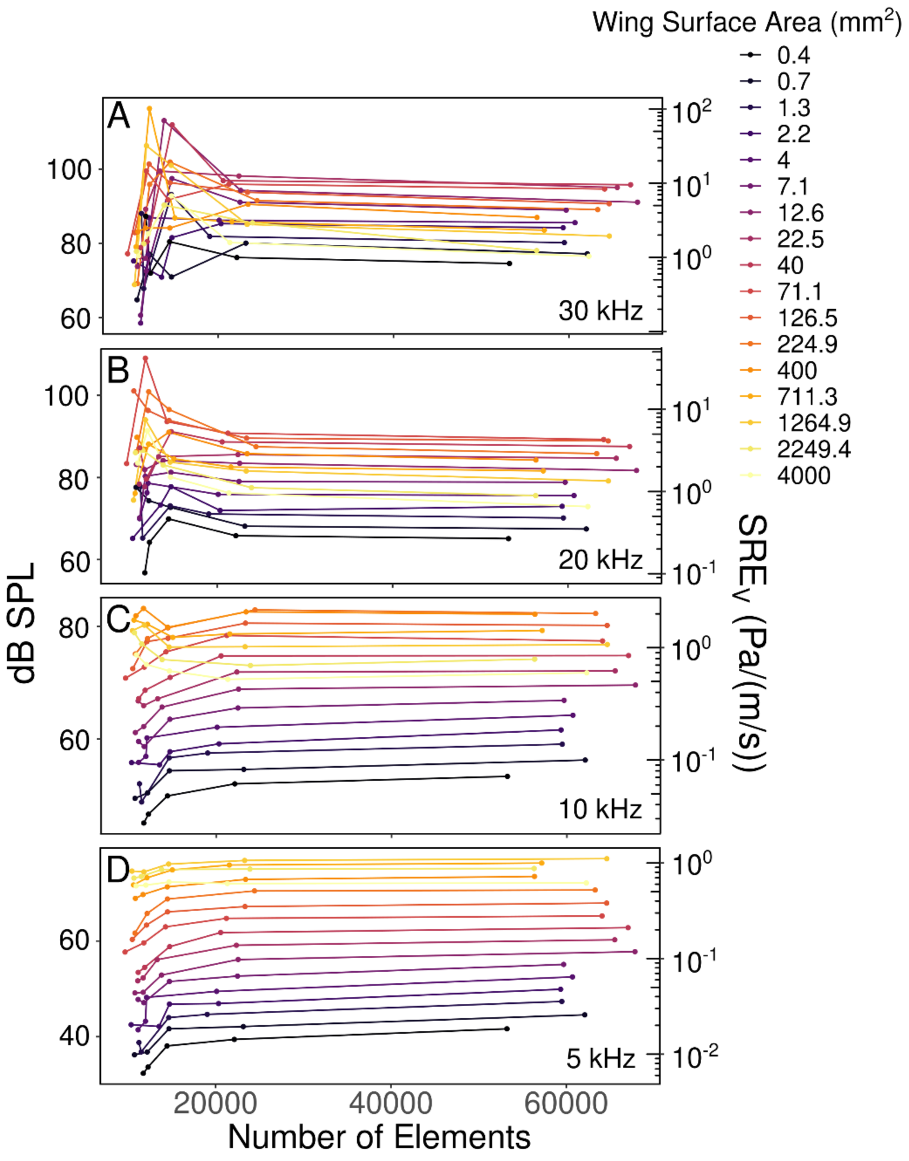


**Figure S5.** Mesh sensitivity analysis for models based on the finite element method. Each line represents the SPL of a single wing size with a different number of mesh elements. Each panel shows this analysis at a different frequency: **A.** 30 kHz, **B.** 20 kHz, **C.** 10 kHz, **D.** 5 kHz. As the difference between the second-largest and largest number of elements was small, we proceeded with the largest number of elements shown here for the analysis.

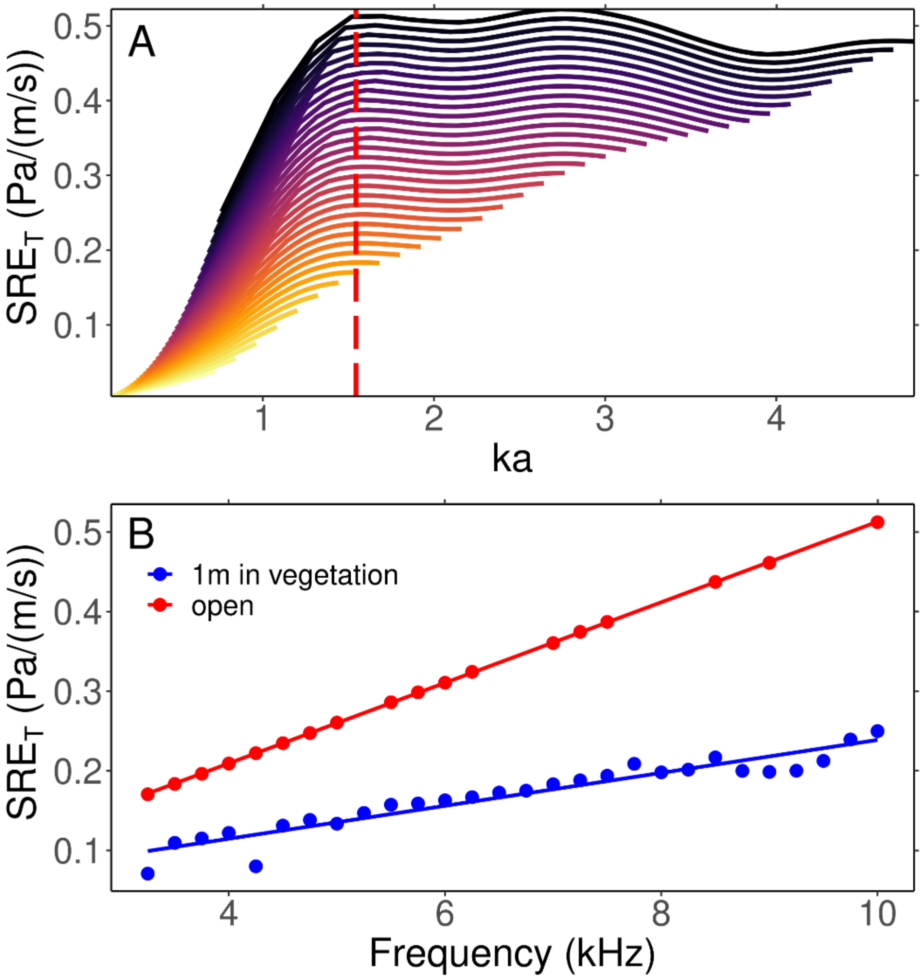


**Figure S6.** Calculating efficiency with optimal baffle. **A.** ka at which optimal efficiency occurs for each frequency. Each frequency is represented by a different line (top line = 10 kHz, bottom line = 1.5 kHz). Line at which efficiency is maximized for each frequency (maximum efficiency ridge) is shown by red dashed line. **B.** Efficiency at this optimal ka (1.55) in open and vegetation conditions.

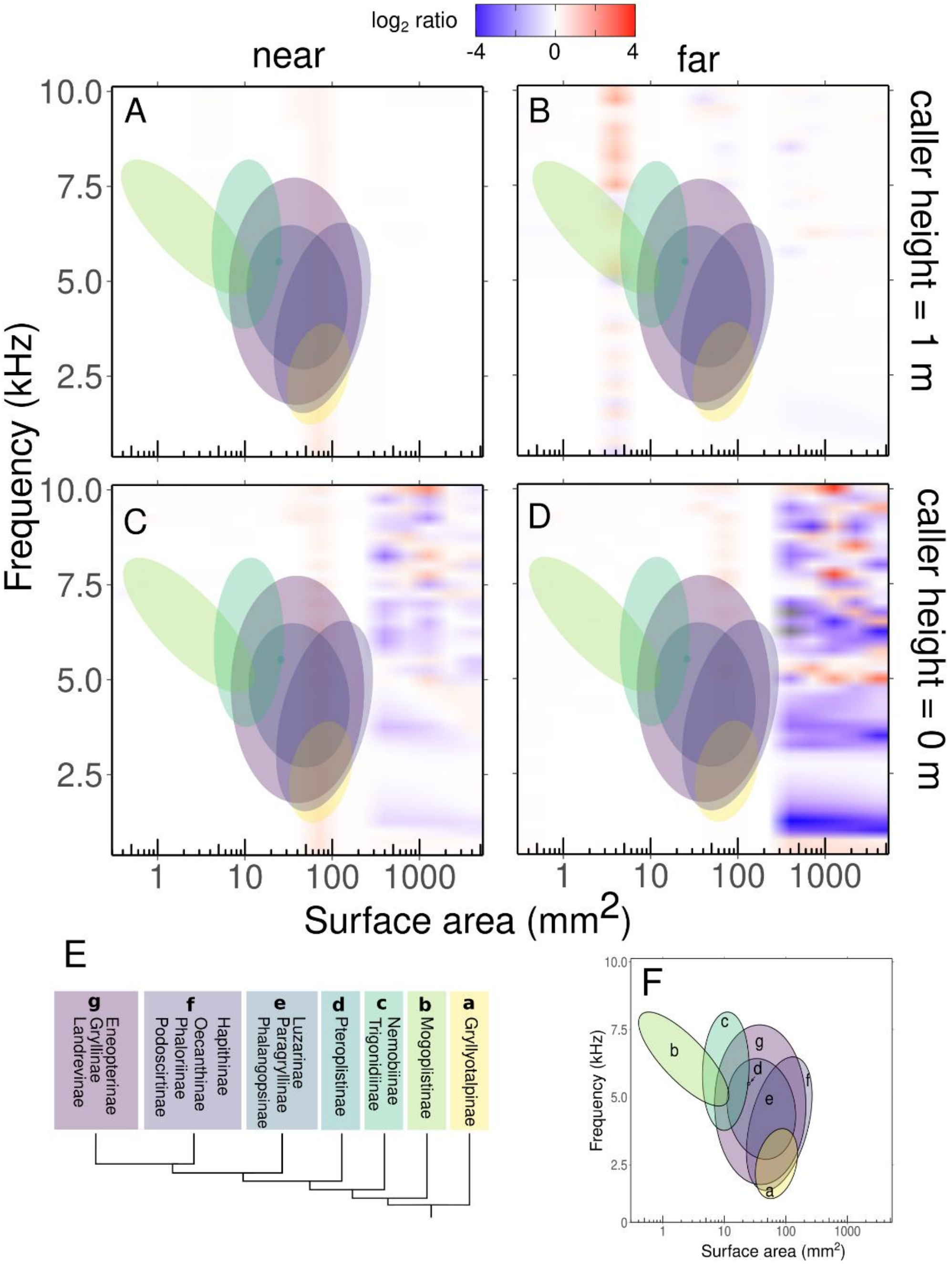


**Figure S7.** Acoustic hardness of ground does not significantly influence $SRE_T$ at biologically relevant ranges of wing size and call frequency. Comparison of $SRE_T$ over hard vs soft ground. Each panel represents a combination of caller height above ground (0 m or 1 m) and receiver distance from caller (0.05 – 02 m "near" and 0.8 – 0.9 m "far"). **A.** Distance = near, height = 1m, **B.** Distance = far, height = 1 m; **C.** Distance = near, height = 1 m, **D.** Distance = far, height = 0 m. Color indicates whether higher $SRE_T$ is found with hard ground (red shades), soft ground (blue shades) or no difference (white). Data are presented as a $\log_2$ ratio instead of a straight proportion. $\text{Log}_2$ ratios are scaled such that the ranges above and below 1 are proportional, rather than values below 1 being compressed between 0 and 1. Each clade of animals is represented by a colored ellipse. **E.** Phylogeny representing each clade **F**. Key to clade represented by each ellipse.

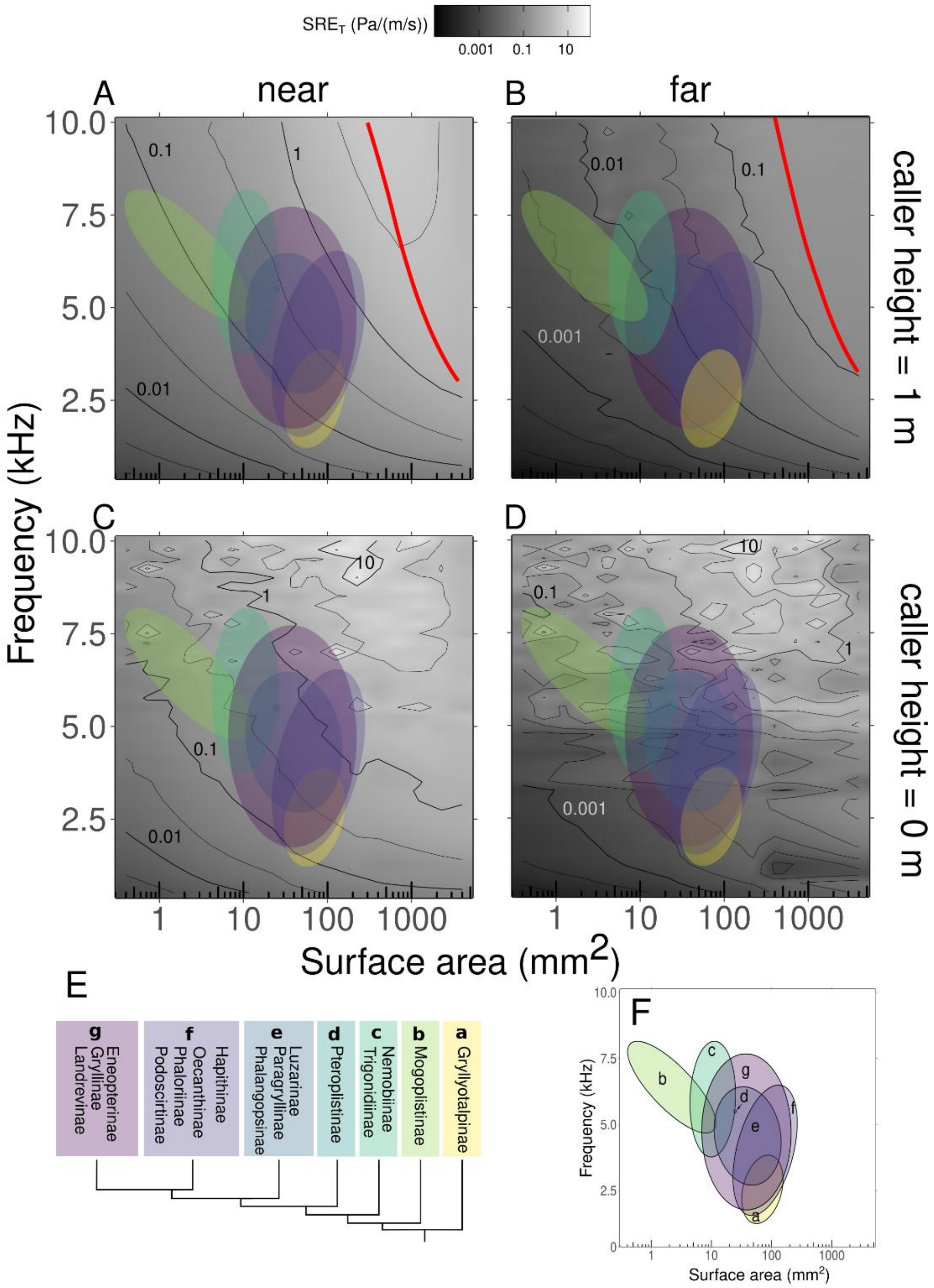


**Figure S8.** Vegetation decreases efficiency overall but does not substantially change the landscape pattern of efficiency. Each panel represents a combination of caller height above ground (0 m or 1 m) and receiver distance from caller (0.05 – 02 m "near" and 0.8 – 0.9 m "far"). In each height and distance scenario, an excess attenuation factor due to vegetation was also applied. **A.** Distance = near, height = 1m, **B.** Distance = far, height = 1 m; **C.** Distance = near, height = 1 m, **D.** Distance = far, height = 0 m. Red lines indicate the radiator size that would minimize short-circuiting for each frequency (an optimally baffled scenario). Each clade of animals is represented by a colored ellipse. **E.** Phylogeny representing each clade **F**. Key to clade represented by each ellipse.

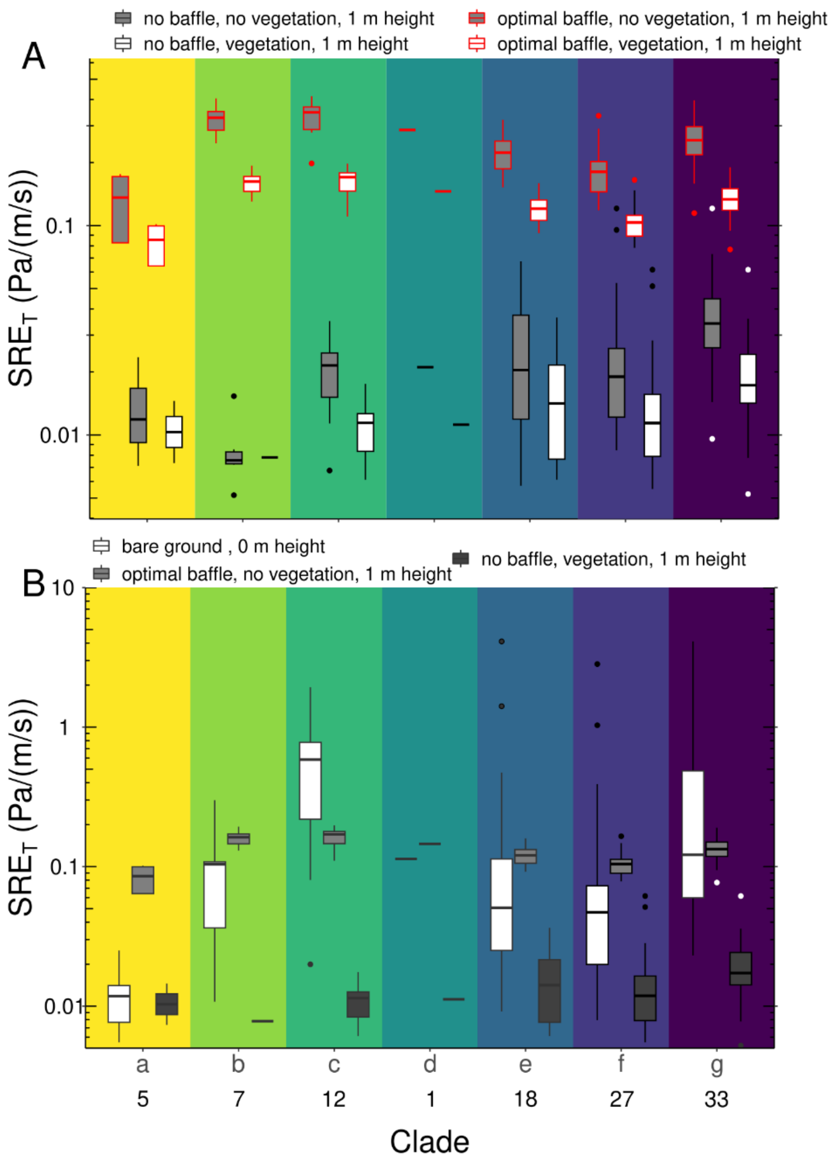


**Figure S9.** The most effective calling strategy (ground vs optimally baffled) varies depending on clade. **A.** Differences in $SRE_T$ by clade depending on vegetation (white bars) and optimal baffle use (grey bars). Vegetation somewhat decreases efficiency in optimally baffled (red outlines) and grounded calling conditions (black outlines). These measurements were taken far from the caller, i.e., an average of the SPL at a distance of 0.8-0.9 m from wings, directly in front of the wings was used. **B.** Comparison of $SRE_T$ in three scenarios, calling near the ground (white bar), or calling from a height from within vegetation with (light grey bar) or without an optimal baffle (dark grey bar). Both strategies, calling from the ground or optimal baffling, offers advantages over calling from a height within vegetation (with the exception of clade a where ground calling seems to perform quite poorly). Species numbers for both panels are given below clade labels.

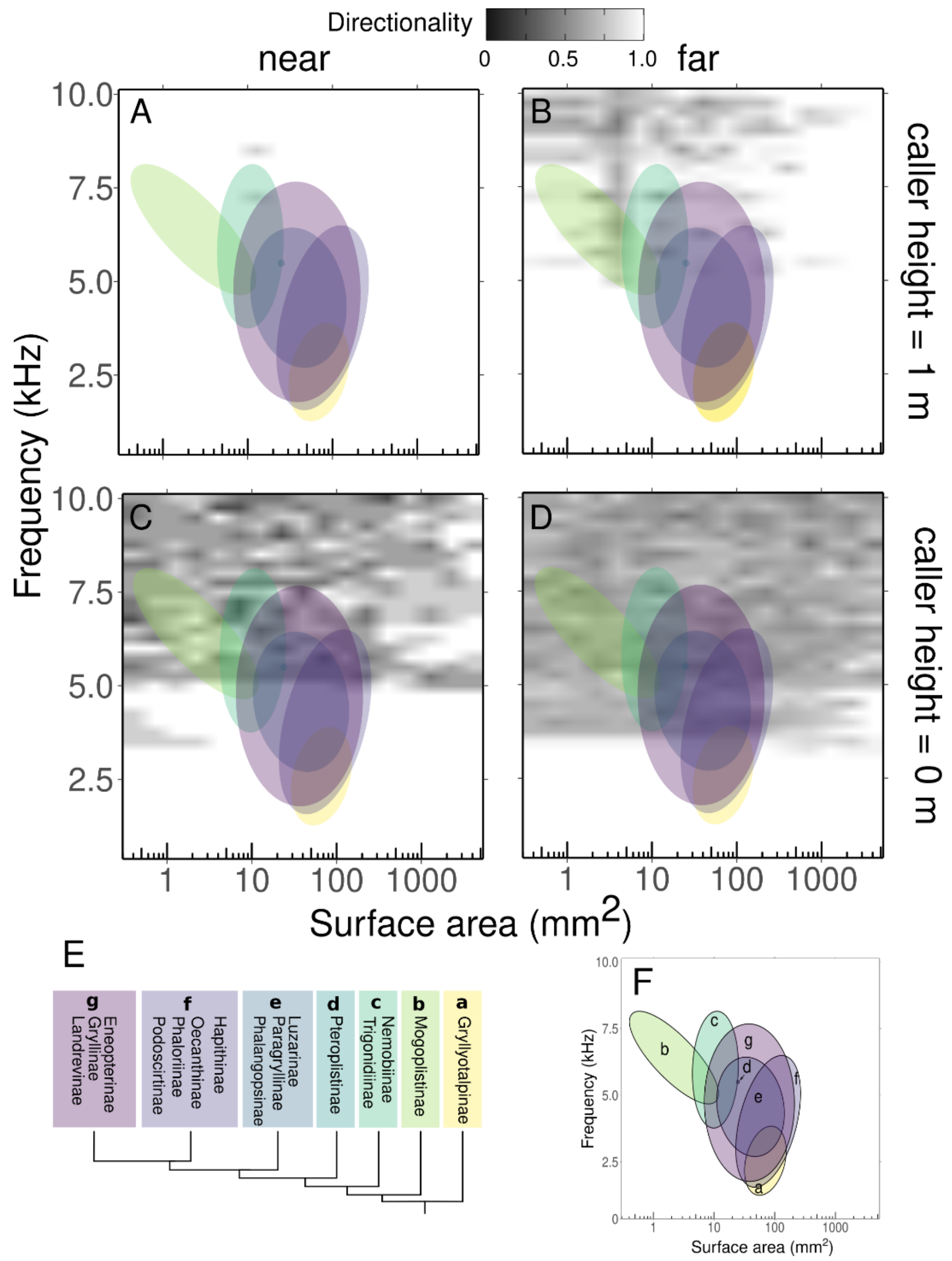


**Figure S10.** Call directionality decreases at higher frequencies, particularly with grounded calling. Each panel represents a combination of caller height above ground (0 m or 1 m) and receiver distance from caller (0.05 – 02 m "near" and 0.8 – 0.9 m "far"). **A.** Distance = near, height = 1m, **B.** Distance = far, height = 1 m; **C.** Distance = near, height = 1 m, **D.** Distance = far, height = 0 m. Each clade of animals is represented by a colored ellipse. **E.** Phylogeny representing each clade **F.** Key to clade represented by each ellipse.

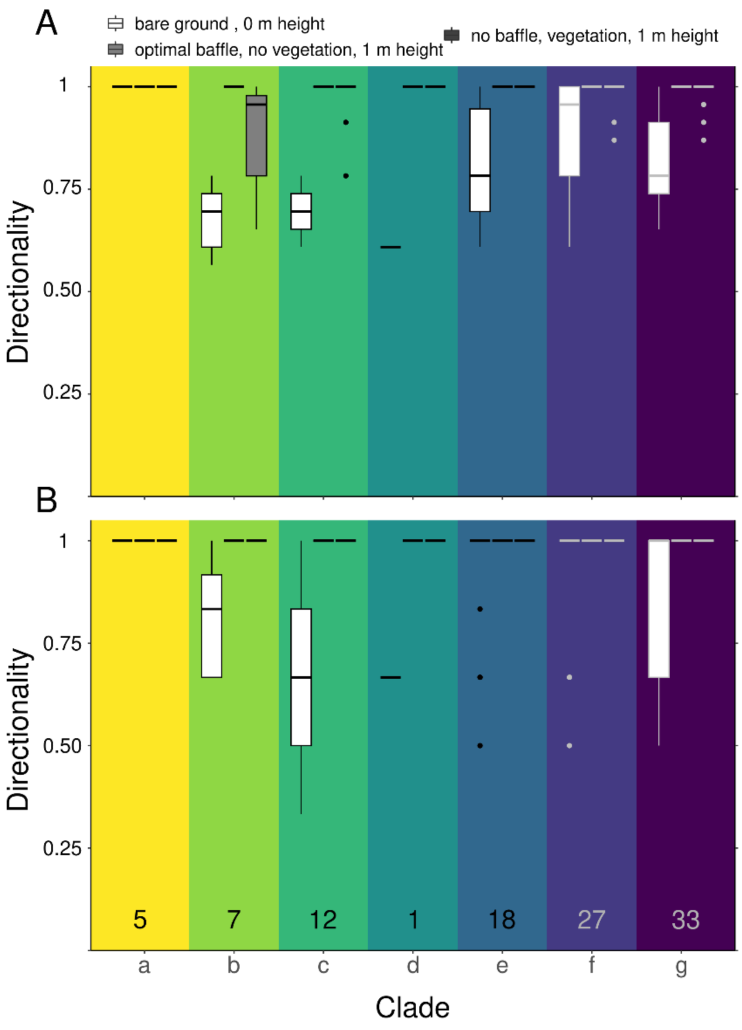


**Figure S11.** Directionality does not vary significantly between different calling strategies. **A.** Directionality in far condition, **B.** Directionality in near condition. Species numbers (both conditions) are indicated above clade labels.

**Table S1**. Sources for morphological data. Asterisk after species name indicates species is represented in both frequency and wing size datasets

| Clade | Subfamily | Genus | Species | Specimen | Relevant reference and/or collection specimen number |
|---|---|---|---|---|---|
| **A** | Gryllotalpinae | *Gryllotalpa* | *australis** | 1 | Orthoptera Species File Specimen ID: 45466 |
| | | | *gryllotalpa** | 1 | Linnean Collection Specimen ID: LINN 8925 |
| | | | *orientalis** | 1 | Orthoptera Species File Taxon ID: 1128860 |
| | | | *permai** | 1 | (Tan and Kamaruddin, 2016) |
| | | | *vineae** | 1 | Museum D'Historie Naturelle ID: MNHN-EO-ENSIF4425 |
| | | | | 2 | Museum D'Historie Naturelle ID: MNHN-EO-ENSIF4425 |
| | | | | 3 | Museum D'Historie Naturelle ID: MNHN-EO-ENSIF4425 |
| **B** | Mogoplistinae | *Cycloptiloides* | *canariensis** | 1 | (Dambach and Gras, 1995) |
| | | *Cycloptilum* | *irregularis** | 1 | (Love and Walker, 1979) |
| | | | *slossoni** | 1 | (Love and Walker, 1979) |
| | | | *tardum** | 1 | (Love and Walker, 1979) |
| | | *Hoplosphyrum* | *boreale** | 1 | (Love and Walker, 1979) |
| | | *Ornebius* | *bimaculatus** | 1 | (Kim, 2011) |
| | | | *kanetataki** | 1 | (Kim, 2011) |
| **C** | Nemobiinae | *Allonemobius* | *allardi** | 1 | Orthoptera Species File Specimen ID: 40715 |
| | | *Bobilla* | *gullanae** | 1 | (Su and Rentz, 2000) |
| | | | *neobivittata** | 1 | (Su and Rentz, 2000) |
| | | *Hygronemobius* | *guriri* | 1 | (Pereira et al., 2013) |
| | | | *indaia** | 1 | (Pereira et al., 2013) |
| | | | *iperoigae** | 1 | (Pereira et al., 2013) |
| | | *Nemobius* | *sylvestris** | 1 | (Barranco et al., 2013) |
| | | *Pteronemobius* | *nigrovus** | 1 | (McIntyre, 1977) |
| | Trigonidiinae | *Anaxipha* | *bradephona** | 1 | Museum D'Historie Naturelle ID: MNHN-EO-ENSIF6482 |

| Clade | Subfamily | Genus | Species | Specimen | Relevant reference and/or collection specimen number |
|---|---|---|---|---|---|
| **C** | Trigonidiinae | *Anaxipha* | *hyalicetra** | 1 | (Cole and Funk, 2019) |
| | | | *tachephona** | 1 | Museum D'Historie Naturelle ID: MNHN-EO-ENSIF6486 |
| | | *Cranistus* | *colliurides** | 1 | (Martins et al., 2012) |
| | | *Phylloscirtus* | *amoenus** | 1 | (Martins et al., 2012) |
| **D** | Pteroplistinae | *Singapuriola* | *separata** | 1 | (Gorochov and Tan, 2012) |
| **E** | Luzarinae | *Lerneca* | *inalata** | 1 | (Lima et al., 2018) |
| | | *Luzaridella* | *susurra** | 1 | (Martins et al., 2013) |
| | | *Vanzoliniella* | *sambophila** | 1 | (Mello and Reis, 1994) |
| | Paragryllinae | *Alcodes* | *chamocoru* | 1 | Orthoptera Species File Specimen ID: 65179 |
| | | | *mococharu* | 1 | Orthoptera Species File Specimen ID: 65181 |
| | | *Aclogryllus* | *crybelos** | 1 | (Nischk and Otte, 2000) |
| | | *Escondacla* | *thymodes** | 1 | Orthoptera Species File Specimen ID: 65198 |
| | | *Neoacla* | *clandestina** | 1 | Orthoptera Species File Specimen ID: 65199 |
| | | *Silvastella* | *epiplatys** | 1 | Orthoptera Species File Specimen ID: 65196 |
| | Phalangopsinae | *Ceyloria* | *latissima* | 1 | Orthoptera Species File Specimen ID: 2983 |
| | | *Endecous* | *betariensis** | 1 | (Mello and Pellegatti-Franco, 1998) |
| | | | *chape** | 1 | (Souza-Dias et al., 2017) |
| | | | *didymus** | 1 | (Desutter-Grandcolas, 2009) |
| | | | *itatibensis** | 1 | (Mello and Pellegatti-Franco, 1998) |
| | | | *naipi* | 1 | (Souza-Dias et al., 2017) |
| | | | *troglobius** | 1 | (Castro-Souza et al., 2020) |
| | | *Lernecella* | *trinitatis* | 1 | Orthoptera Species File Taxon ID: 1125930 |
| | | *Pseudotrigonidium* | *personatum* | 1 | (Desutter-Grandcolas, 2009) |

| Clade | Subfamily | Genus | Species | Specimen | Relevant reference and/or collection specimen number |
|---|---|---|---|---|---|
| **E** | Phalangopsinae | *Tremellia* | *timah** | 1 | (Gorochov and Tan, 2012) |
| | Phaloriinae | *Phaloria* | *anapina** | 1 | (Otte, 2007) |
| | | | *chopardi** | 1 | (Desutter-Grandcolas, 2009) |
| | | | *jerelynae** | 1 | (Gorochov and Tan, 2012) |
| | | *Trellius* | *neesoon* | 1 | (Gorochov and Tan, 2012) |
| **F** | Hapithinae | *Hapithus* | *agitator** | 1 | Orthoptera Species File Specimen ID: 138599 |
| | | | *vagus** | 1 | Orthoptera Species File Specimen ID: 65035 |
| | Oecanthinae | *Neoxabea* | *bipunctata** | 1 | Image captured for present study in lab |
| | | | *brevipes** | 1 | (Zefa et al., 2018) |
| | | | *cerrojesusensis** | 1 | Image captured for present study in lab |
| | | | *ottei** | 1 | Image captured for present study in lab |
| | | *Oecanthus* | *alexanderi** | 1 | Image captured for present study in lab |
| | | | *angustus** | 1 | PaDILspecies ID: *Oecanthus angustus* |
| | | | *argentinus** | 1 | University of British Columbia Insect Collection, Specimen: SEM-UBC GRY-0951 |
| | | | *forbsei** | 1 | Image captured for present study in lab |
| | | | *fultoni** | 1 | Orthoptera Species File Specimen ID: 40710 |
| | | | *henryi** | 1 | (Metrani and Balakrishnan, 2005) |
| | | | *latipennis** | 1 | University of Guelph Insect Collection: Specimen BIOUG44550-E07 |
| | | | | 2 | University of Guelph Insect Collection: Specimen BIOUG44550-E08 |
| | | | *lineolatus** | 1 | (Zefa et al., 2012) |
| | | | *major* | 1 | Orthoptera Species File Specimen ID: 40712 |
| | | | *nigricornis** | 1 | Orthoptera Species File Taxon ID: 345166 |
| | | | *niveus** | 1 | Orthoptera Species File Taxon ID: 345151 |

| Clade | Subfamily | Genus | Species | Specimen | Relevant reference and/or collection specimen number |
|---|---|---|---|---|---|
| **F** | Oecanthinae | *Oecanthus* | *pallidus** | 1 | (Zefa et al., 2012) |
| | | | *pictus** | 1 | (Milach et al., 2015) |
| | | | *pini** | 1 | Image captured for present study in lab |
| | | | *quadripunctatus** | 1 | https://www.insectimages.org/browse/subthumb.cfm?sub=9113 |
| | | | | 2 | UBC Database ID: SEM-UBC GRY-0918 |
| | | | *rileyi** | 1 | Orthoptera Species File Taxon ID: 1128127 |
| | | | *rufescens** | 1 | NHM Specimen ID: 012497644 |
| | | | | 2 | NHM Specimen ID: 012497645 |
| | | | | 3 | NHM Specimen ID: 012497646 |
| | | | | 4 | PaDIL species ID: *Oecanthus rufescens* |
| | | | *texensis** | 1 | Image captured for present study in lab |
| | | | *valensis* | 1 | (Milach et al., 2016) |
| | | | *varicornis** | 1 | Image captured for present study in lab |
| | Podoscirtinae | *Madasumma* | *affinis** | 1 | (Otte, 2007) |
| | | *Truljalia* | *formosa** | 1 | (He, 2012) |
| | Podoscirtinae | *Varitrella* | *suikei** | 1 | (Tan et al., 2020) |
| **G** | Eneopterinae | *Agnotecous* | *azurensis** | 1 | (Desutter-Grandcolas and Robillard, 2006) |
| | | | *brachypterus** | 1 | (Robillard et al., 2010) |
| | | | *meridionalis** | 1 | Museum D’Historie Naturelle ID: MNHN-EO-ENSIF1775 |
| | | | *pinsula** | 1 | (Robillard et al., 2010) |
| | | | *sarramea** | 1 | Museum D’Historie Naturelle ID: MNHN-EO-ENSIF988 |
| | | | *yahoue** | 1 | (Desutter-Grandcolas and Robillard, 2006) |
| | | *Arilpa* | *binderia** | 1 | (Otte, 2007) |

| Clade | Subfamily | Genus | Species | Specimen | Relevant reference and/or collection specimen number |
|---|---|---|---|---|---|
| **G** | Eneopterinae | *Arilpa* | *gidya** | 1 | (Otte, 2007) |
| | | *Cardiodactylus* | *guttulus** | 1 | (Robillard and Ichikawa, 2009) |
| | | | *novaeguinea** | 1 | (Robillard and Ichikawa, 2009) |
| | | *Eurepa* | *bifasciata** | 1 | (Robillard and Su, 2018) |
| | | *Gnominthus* | *baitabagus** | 1 | (Robillard and Su, 2018) |
| | | *Lebinthus* | *bitaeniatus** | 1 | (Robillard et al., 2013) |
| | | | *luae** | 1 | Museum D'Historie Naturelle ID: MNHN-EO-ENSIF3208 |
| | | *Myara* | *pakaria** | 1 | (Otte, 2007) |
| | | | *wintrena** | 1 | (Otte, 2007) |
| | | *Pixibinthus* | *sonicus** | 1 | (Anso et al., 2016) |
| | | *Salmanites* | *peekara** | 1 | (Otte, 2007) |
| | | *Xenogryllus* | *eneopteroides** | 1 | (Jaiswara et al., 2019) |
| | | | *transversus** | 1 | (Jaiswara et al., 2019) |
| | Gryllinae | *Eurepella* | *mjobergi** | 1 | PaDIL species ID: *Eurepella mjobergi* |
| | | *Gryllus* | *amarensis* | 1 | Museum D'Historie Naturelle ID: 7031 |
| | | | *assimilis** | 1 | SINA species ID: *Gryllus assimilis* |
| | | | *bimaculatus** | 1 | Orthoptera Species File Taxon ID: 1122377 |
| | | | *brevicaudus** | 1 | SINA species ID: *Gryllus brevicaudus* |
| | | | *campestris** | 1 | Need to figure out specific specimen |
| | | | | 2 | Need to figure out specific specimen |
| | | | *carvalhoi* | 1 | Museum D'Historie Naturelle ID: MNHN-EO-ENSIF7242 |
| | | | *chaldeus* | 1 | Museum D'Historie Naturelle ID: MNHN-EO-ENSIF7192 |
| | | | *chappuisi** | 1 | Museum D'Historie Naturelle ID: MNHN-EO-ENSIF7046 |

| Clade | Subfamily | Genus | Species | Specimen | Relevant reference and/or collection specimen number |
|---|---|---|---|---|---|
| **G** | Gryllinae | *Gryllus* | *cohni** | 1 | (Weissman and Gray, 2019) |
| | | | *firmus** | 1 | (Weissman and Gray, 2019) |
| | | | *fultoni** | 1 | Orthoptera Species File Specimen ID: 40672 |
| | | | *lineaticeps** | 1 | (Weissman and Gray, 2019) |
| | | | *multipulsator** | 1 | (Weissman and Gray, 2019) |
| | | | *pennsylvanicus** | 2 | Orthoptera Species File Specimen ID: 43773 |
| | | | | 3 | UBC Database ID: SEM-UBC GRY-0542 |
| | | | *veletis** | 1 | Orthoptera Species File Specimen ID: 40674 |
| | | | | 2 | UBC Database ID: SEM-UBC GRY-0643 |
| | | | *vocalis** | 1 | Orthoptera Species File Specimen ID: 64224 |
| | | *Miogryllus* | *itaquiensis** | 1 | (Orsini et al., 2017) |
| | | | *piracicabensis** | 1 | (Orsini et al., 2017) |
| | | *Teleogryllus* | *commodus** | 1 | (Otte, 2007) |
| | | | *marini** | 1 | (Otte, 2007) |
| | | | *oceanicus** | 1 | (Otte, 2007) |
| | Itarinae | *Itara* | *kirejtshuki** | 1 | NMHUK 012497661 |
| | | | *minor** | 1 | Museum D'Historie Naturelle ID: MNHN-EO-ENSIF8162 |
| | Landrevinae | *Striduleva* | *crepitans** | 1 | Museum D'Historie Naturelle ID: MNHN-EO-ENSIF2059 |

**Table S2.** Sources for call frequency data. Asterisk after species name indicates species is represented in both frequency and wing size datasets

| Clade | Subfamily | Genus | Species | Recording | Relevant reference and/or collection specimen number |
|---|---|---|---|---|---|
| **A** | Gryllotalpa | *Gryllotalpa* | *australis** | 1-47 | (Kavanagh and Young, 1989) (range of values given in publication) |
| | | | *fulvipes* | 1 | (Tan and Kamaruddin, 2016) |
| | | | *gryllotalpa** | 1 | Orthoptera Species File Sound ID: 1176 |
| | | | *permai** | 1 | (Tan and Kamaruddin, 2016) |
| | | | *vineae** | 1 | Orthoptera Species File Sound ID: 1198 |
| | | | *canariensis** | 1 | (Dambach and Gras, 1995) |
| **B** | Mogoplistinae | *Cycloptiloides* | *irregularis** | 1 | Crickets north of Mexico species Id: Key's scaly cricket |
| | | *Cycloptilum* | *slossoni** | 1 | Crickets north of Mexico species Id: Slosson's scaly cricket |
| | | | *tardum** | 1 | (Otte, 2007) |
| | | | *boreale** | 1 | Crickets north of Mexico species Id: long-winged scaly cricket |
| | | *Hoplosphyrum* | *bimaculatus** | 1 | (He et al., 2017) |
| | | *Ornebius* | *kanetataki** | 1 | (He et al., 2017) |
| | | | *allardi** | 1 | Crickets north of Mexico species Id: Allard's ground cricket |
| **C** | Nemobiinae | *Allonemobius* | *gullanae** | 1 | (Su and Rentz, 2000) |
| | | *Bobilla* | *neobivittata** | 1 | (Su and Rentz, 2000) |
| | | | *indaia** | 1 | (Pereira et al., 2013) |
| | | *Hygronemobius* | *iperoigae** | 1 | (Pereira et al., 2013) |
| | | | *sylvestris** | 1 | Orthoptera Species File Sound ID: 1045 |

| Clade | Subfamily | Genus | Species | Recording | Relevant reference and/or collection specimen number |
|---|---|---|---|---|---|
| **C** | Nemobiinae | *Nemobius* | *nigrovus** | 1 | (McIntyre, 1977) |
| | | *Pteronemobius* | *bradephona** | 1 | Orthoptera Species File Sound ID: 1832 |
| | Trigonidiinae | *Anaxipha* | *hyalicetra** | 1 | (Cole and Funk, 2019) |
| | | | *tachephona** | 1 | Orthoptera Species File Sound ID: 1833 |
| | | | *colliurides** | 1 | (Martins et al., 2012) |
| | | *Cranistus* | *amoenus** | 1 | (Martins et al., 2012) |
| | | *Phylloscirtus* | *separata** | 1 | (Gorochov and Tan, 2012) |
| **D** | Pteroplistinae | *Singapuriola* | *inalata** | 1 | (Lima et al., 2018) |
| **E** | Luzarinae | *Lerneca* | *susurra** | 1 | (Martins et al., 2013) |
| | | *Luzaridella* | *sambophila** | 1 | (Mello and Reis, 1994) |
| | | *Vanzoliniella* | *chamocoru** | 1 | (Nischk and Otte, 2000) |
| | Paragryllinae | *Aclodes* | *moccharacharu** | 1 | (Nischk and Otte, 2000) |
| | | | *crybelos** | 1 | (Nischk and Otte, 2000) |
| | | *Aclogryllus* | *thymodes** | 1 | (Nischk and Otte, 2000) |
| | | *Escondacla* | *clandestine** | 1 | (Nischk and Otte, 2000) |
| | | *Neoacla* | *epiplatys** | 1 | (Nischk and Otte, 2000) |
| | | *Silvastella* | *betariensis** | 1 | (He, 2012) |
| | Phalangopsinae | *Endecous* | *chape** | 1 | (Souza-Dias et al., 2017) |

| **Clade** | **Subfamily** | **Genus** | **Species** | **Recording** | **Relevant reference and/or collection specimen number** |
|---|---|---|---|---|---|
| **E** | Phalangopsinae | *Endecous* | *didymus** | 1 | (Castro-Souza et al., 2020) |
| | | | *itatibensis** | 1 | (Mello and Pellegatti-Franco, 1998) |
| | | | *troglobius** | 1 | (Castro-Souza et al., 2020) |
| | | | *timah** | 1 | (Gorochov and Tan, 2012) |
| | | *Tremellia* | *anapina** | 1 | (Su and Rentz, 2000) |
| | Phaloriinae | *Phaloria* | *chopardi** | 1 | (Desutter-Grandcolas, 2009) |
| | | | *jerelynae** | 1 | (Gorochov and Tan, 2012) |
| | | | *baitabagus** | 1 | (Vicente et al., 2015) |
| **F** | Hapithinae | *Hapithus* | *melodius* | 1 | Handbook of crickets and katydids |
| | | | *vagus** | 1 | Macaulay Library asset: 114470 |
| | | | *diplastes* | 1 | Handbook of crickets and katydids |
| | | *Orocharis* | *gryllodes* | 1 | Handbook of crickets and katydids |
| | | | *luteolira* | 1 | Handbook of crickets and katydids |
| | | | *nigrifrons* | 1 | Handbook of crickets and katydids |
| | | | *saltator* | 1 | Handbook of crickets and katydids |
| | | | *tricornis* | 1 | Handbook of crickets and katydids |
| | | | *bipunctata** | 1 | Crickets north of Mexico species ID: *Neoxabea bipunctata* |
| | Oecanthinae | *Neoxabea* | *brevipes ** | 1 | (Zefa et al., 2018) |

| **Clade** | **Subfamily** | **Genus** | **Species** | **Recording** | **Relevant reference and/or collection specimen number** |
|---|---|---|---|---|---|
| **F** | *Oecanthinae* | *Oecanthus* | *cerrojesusensis** | 1 | Orthoptera Species File Sound ID: 2345 |
| | | | *ottei** | 1 | Orthoptera Species File Sound ID: 2346 |
| | | | *alexanderi** | 1 | Crickets North of Mexico Species ID: *Oecanthus alexanderi* |
| | | | *angustus** | 1 | (Otte, 2007) |
| | | | *argentinus** | 1 | Crickets North of Mexico Species ID: *Oecanthus argentinus* |
| | | | *californicus* | 2 | Orthoptera Species File Sound ID: 1535 |
| | | | | 1 | Crickets North of Mexico Species ID: *Oecanthus californicus* |
| | | | *forbsei** | 1 | Crickets North of Mexico Species ID: *Oecanthus forbsei* |
| | | | *fultoni** | 1 | Crickets North of Mexico Species ID: *Oecanthus fultoni* |
| | | | *henryi** | 1 | (Metrani and Balakrishnan, 2005) |
| | | | *latipennis** | 1 | Crickets North of Mexico Species ID: *Oecanthus latipennis* |
| | | | | 2 | Orthoptera Species File Sound ID: 1002 |
| | | | *lineolatus** | 1 | (Zefa et al., 2012) |
| | | | *nigricornis** | 1 | Crickets North of Mexico Species ID: *Oecanthus nigricornis* |
| | | | *niveus** | 1 | Crickets North of Mexico Species ID: *Oecanthus niveus* |
| | | | *pallidus** | 1 | (Zefa et al., 2012) |
| | | | *pictus** | 1-9 | Orthoptera Species File Taxon ID: 1223417 (9 songs from different temperatures) |
| | | | *pini** | 1 | Crickets North of Mexico Sound File: 587sl |

| Clade | Subfamily | Genus | Species | Recording | Relevant reference and/or collection specimen number |
|---|---|---|---|---|---|
| **F** | Oecanthinae | *Oecanthus* | *quadripunctatus** | 1 | Orthoptera Species File Sound File 1531 |
| | | | *rileyi** | 1 | Orthoptera Species File Sound File: 1540 |
| | | | *rufescens** | 1 | (Otte, 2007) |
| | | | *texensis** | 1 | (Symes and Collins, 2013) |
| | | | *varicornis** | 1 | Crickets North of Mexico Sound File: 593sl |
| | | | *walker* | 1 | Crickets North of Mexico Species ID: *Oecanthus walkeri* |
| | | | *affinis** | 1 | (Otte, 2007) |
| | Podoscirtinae | *Madasumma* | *jirranda* | 1 | (Otte, 2007) |
| | | | *kanina* | 1 | (Otte, 2007) |
| | | | *loorea* | 1 | (Otte, 2007) |
| | | | *formosa** | 1 | (He, 2012) |
| | | *Truljalia* | *suikei** | 1 | (Tan et al., 2020) |
| | | *Varitrella* | *azurensis** | 1 | Museum D'Historie Naturelle ID: MNHN-SO-2018-100 |
| **G** | Eneopterinae | *Agnotecous* | *brachypterus** | 1 | (Robillard et al., 2010) |
| | | | *clarus* | 1 | Museum D'Historie Naturelle ID: MNHN-SO-2018-102 |
| | | | *meridionalis** | 1 | Museum D'Historie Naturelle ID: MNHN-SO-2018-99 |
| | | *Agnotecous* | *pinsula** | 1 | (Robillard et al., 2010) |
| | | | *sarramea** | 1 | (Robillard and Desutter-Grandcolas, 2004) |

| Clade | Subfamily | Genus | Species | Recording | Relevant reference and/or collection specimen number |
|---|---|---|---|---|---|
| **G** | Eneopterinae | *Agnotecous* | *yahoue** | 1 | (Robillard and Desutter-Grandcolas, 2004) |
| | | | *binderia** | 1 | (Otte, 2007) |
| | | *Arilpa* | *gidya** | 1 | (Otte, 2007) |
| | | | *wirrilla* | 1 | (Otte, 2007) |
| | | | *guttulus** | 1 | (Robillard and Ichikawa, 2009) |
| | | *Cardiodactylus* | *novaeguinea** | 1 | (Otte, 2007) |
| | | | *bifasciata** | 1 | (Otte, 2007) |
| | | *Eurepa* | *eeboolaga* | 1 | (Otte, 2007) |
| | | | *marginipennis* | 1 | (Otte, 2007) |
| | | | *noarana* | 1 | (Otte, 2007) |
| | | | *nurndina* | 1 | (Otte, 2007) |
| | | | *wirkutta* | 1-2 | (Otte, 2007) (range of values given in publication) |
| | | | *woortooa* | 1 | (Otte, 2007) |
| | | | *yumbena* | 1 | (Otte, 2007) |
| | | | *bitaeniatus** | 1 | (Robillard and Tan, 2013) |
| | | *Gnominthus* | *baitabagus** | 1 | (Anso et al., 2016) |
| | | *Lebinthus* | *luae** | 1 | (Robillard and Tan, 2013) |
| | | | *aperta* | 1 | (Otte, 2007) |

| **Clade** | **Subfamily** | **Genus** | **Species** | **Recording** | **Relevant reference and/or collection specimen number** |
|---|---|---|---|---|---|
| **G** | Eneopterinae | *Myara* | *marimbula* | 1 | (Otte, 2007) |
| | | | *muttaburra* | 1 | (Otte, 2007) |
| | | | *pakaria** | 1 | (Otte, 2007) |
| | | | *sordida* | 1 | (Otte, 2007) |
| | | | *unicolor* | 1-2 | (Otte, 2007) (range of values given in publication) |
| | | | *wintrena** | 1 | (Robillard and Desutter-Grandcolas, 2004) |
| | | | *yurgama* | 1 | (Otte, 2007) |
| | | | *vittatus* | 1 | (Robillard and Desutter-Grandcolas, 2004) |
| | | *Nisitrus* | *allaris* | 1 | (Otte, 2007) |
| | | *Pixibinthus* | *sonicus** | 1 | (Anso et al., 2016) |
| | | *Salmanites* | *ninbella* | 1 | (Otte, 2007) |
| | | | *noccundris* | 1 | (Otte, 2007) |
| | | | *noonamina* | 1 | (Otte, 2007) |
| | | | *peekara** | 1 | (Otte, 2007) |
| | | | *poene* | 1 | (Otte, 2007) |
| | | | *taltantris* | 1 | (Otte, 2007) |
| | | | *terba* | 1-2 | (Otte, 2007) (range of values given in publication) |
| | | | *wittilliko* | 1 | (Otte, 2007) |

| Clade | Subfamily | Genus | Species | Recording | Relevant reference and/or collection specimen number |
|---|---|---|---|---|---|
| **G** | Eneopterinae | *Salmanites* | *eneopteroides** | 1 | (Jaiswara et al., 2019) |
| | | *Xenogryllus* | *maichauensis* | 1 | (Jaiswara et al., 2019) |
| | | | *marmoratus* | 1 | (Jaiswara et al., 2019) |
| | | | *mozambicus* | 1 | (Jaiswara et al., 2019) |
| | | | *transversus** | 1 | Database found within http://www.biologie.uni-ulm.de, no longer exists |
| | | | | 2 | Database found within http://www.biologie.uni-ulm.de, no longer exists |
| | | | *ululiu* | 1 | Database found within http://www.biologie.uni-ulm.de, no longer exists |
| | | | *ballina* | 1 | (Otte, 2007) |
| | Gryllinae | *Eurepella* | *iando* | 1 | (Otte, 2007) |
| | | | *jillangolo* | 1 | (Otte, 2007) |
| | | | *kulkawirra* | 1 | (Otte, 2007) |
| | | | *lewara* | 1 | (Otte, 2007) |
| | | | *mataranka* | 1 | (Otte, 2007) |
| | | | *meda* | 1 | (Otte, 2007) |
| | | | *mjobergi** | 1-2 | (Otte, 2007) (range of values given in publication) |
| | | | *moojerra* | 1 | (Otte, 2007) |
| | | | *oana* | 1 | (Otte, 2007) |
| | | | *quarriana* | 1 | (Otte, 2007) |

| Clade | Subfamily | Genus | Species | Recording | Relevant reference and/or collection specimen number |
|---|---|---|---|---|---|
| **G** | Gryllinae | *Eurepella* | *tinga* | 1 | (Otte, 2007) |
| | | | *tjairaia* | 1 | (Otte, 2007) |
| | | | *torowatta* | 1 | (Otte, 2007) |
| | | | *wanga* | 1 | (Otte, 2007) |
| | | | *waninga* | 1 | (Otte, 2007) |
| | | *Gryllus* | *assimilis** | 1 | Crickets North of Mexico sound file: 483sl |
| | | | | 2 | Crickets North of Mexico sound file: 483ss2 |
| | | | *bimaculatus** | 1 | Orthoptera Species File Sound ID: 1295 |
| | | | *brevicaudus** | 1 | Crickets North of Mexico sound file: 465sldw |
| | | | | 2 | Crickets North of Mexico sound file: 465ss2wg |
| | | | *campestris** | 1 | Orthoptera Species File sound ID: 1741 |
| | | | *chappuisi** | 1 | Orthoptera Species File sound ID: 1739 |
| | | | *cohni** | 1 | Crickets North of Mexico sound file: 722sl |
| | | | *firmus** | 1 | Crickets North of Mexico sound file: 481sl |
| | | | *fultoni** | 1 | Crickets North of Mexico sound file: 484sl |
| | | | | 2 | Crickets North of Mexico sound file: 484slc |
| | | | *lineaticeps** | 1 | Crickets North of Mexico sound file: 467sldw |
| | | | *multipulsator** | 1 | Crickets North of Mexico sound file: 499sl |

| Clade | Subfamily | Genus | Species | Recording | Relevant reference and/or collection specimen number |
|---|---|---|---|---|---|
| **G** | Gryllinae | *Gryllus* | *multipulsator** | 2 | Crickets North of Mexico sound file: 499slwg |
| | | | *pennsylvanicus** | 1 | Orthoptera Species File sound ID: 1258 |
| | | | *texensis* | 1 | Crickets North of Mexico sound file: 479sl |
| | | | *veletis** | 1 | Crickets North of Mexico sound file: 488sl |
| | | | *vocalis** | 1 | Crickets North of Mexico sound file: 466sldw |
| | | | *itaquiensis** | 1 | (Otte, 2007) |
| | | *Miogryllus* | *piracicabensis** | 1-30 | (Orsini et al., 2017) (range of values given in publication) |
| | | | *commodus** | 1 | (Otte, 2007) |
| | | *Teleogryllus* | *marini** | 1-2 | (Otte, 2007) (range of values given in publication) |
| | | | *oceanicus** | 1 | (Otte, 2007) |
| | | *Itara* | *kirejtshuki** | 1 | Orthoptera Species File sound ID: 1796 |
| | | | *minor** | 1 | Database found within http://www.biologie.uni-ulm.de, no longer exists |
| | | *Striduleva* | *crepitans** | 1-2 | (Hugel, 2009) (range of values given in publication) |

Full citations for Specimen data:

Anso J, Barrabé L, Desutter-Grandcolas L, Jourdan H, Grandcolas P, Dong J, Robillard T. 2016. Old Lineage on an Old Island: Pixibinthus, a New Cricket Genus Endemic to New Caledonia Shed Light on Gryllid Diversification in a Hotspot of Biodiversity. *PLOS ONE* **11**:e0150920. doi:10.1371/journal.pone.0150920

Barranco P, Gilgado JD, Ortuño VM. 2013. A new mute species of the genus Nemobius Serville (Orthoptera, Gryllidae, Nemobiinae) discovered in colluvial, stony debris in the Iberian Peninsula: A biological, phenological and biometric study. *Zootaxa* **3691**:201–219. doi:10.11646/zootaxa.3691.2.1

Castro-Souza RA, Zefa E, Ferreira RL. 2020. New troglobitic and troglophilic syntopic species of Endecous (Orthoptera, Grylloidea, Phalangopsidae) from a Brazilian cave: a case of sympatric speciation? *Zootaxa* **4810**:271–304. doi:10.11646/zootaxa.4810.2.3

Cole JA, Funk DH. 2019. Anaxipha hyalicetra sp. n. (Gryllidae: Trigonidiinae), a new sword-tailed cricket species from Arizona. *Journal of Orthoptera Research* **28**:3–9.

Dambach M, Gras A. 1995. Bioacoustics of a miniature cricket, Cycloptiloides canariensis (Orthoptera: Gryllidae: Mogoplistinae). *Journal of Experimental Biology* **198**:721–728. doi:10.1242/jeb.198.3.721

Desutter-Grandcolas L. 2009. New and little known crickets from Espiritu Santo Island, Vanuatu (Insecta, Orthoptera, Grylloidea, Pseudotrigonidium Chopard, 1915, Phaloriinae and Nemobiinae p.p.). *zoos* **31**:619–659. doi:10.5252/z2009n3a12

Desutter-Grandcolas L, Robillard T. 2006. Phylogenetic systematics and evolution of Agnotecous in New Caledonia (Orthoptera: Grylloidea, Eneopteridae). *Systematic Entomology* **31**:65–92. doi:10.1111/j.1365-3113.2005.00299.x

Gorochov AV, Tan MK. 2012. New crickets of the subfamilies Phaloriinae and Pteroplistinae (Orthoptera: Gryllidae) from Singapore. *Zootaxa* **3525**:18–34. doi:10.11646/zootaxa.3525.1.2

He Z. 2012. A new species of Truljalia Gorochov 1985 from Taiwan (Orthoptera: Gryllidae; Podoscirtinae; Podoscirtini). *Zootaxa* **3591**:79–83. doi:10.11646/zootaxa.3591.1.5

He Z, Lu H, Liu Y, Li K. 2017. A new species of Ornebius Guérin-Méneville, 1844 from East China (Orthoptera: Mogoplistidae: Mogoplistinae). *Zootaxa* **4303**:445–450. doi:10.11646/zootaxa.4303.3.10

Hugel S. 2009. New Landrevinae from Mascarene islands and little known Landrevinae from Africa and Comoros (Grylloidea: Landrevinae). *Annales de la Société entomologique de France (NS)* **45**:193–215. doi:10.1080/00379271.2009.10697602

Jaiswara R, Dong J, Ma L, Yin H, Robillard T. 2019. Taxonomic revision of the genus Xenogryllus Bolívar, 1890 (Orthoptera, Gryllidae, Eneopterinae, Xenogryllini). *Zootaxa* **4545**:301–338. doi:10.11646/zootaxa.4545.3.1

Kavanagh MW, Young D. 1989. Bilateral symmetry of sound production in the mole cricket, Gryllotalpa australis. *J Comp Physiol A* **166**:43–49. doi:10.1007/BF00190208

Kim T. 2011. A Taxonomic Study of the Scaly Cricket Family Mogoplistidae (Orthoptera: Ensifera: Grylloidea) in Korea. *Zootaxa* **2928**:41–48. doi:10.11646/zootaxa.2928.1.4

Lima RM, Schuchmann K-L, Tissiani AS, Nunes LA, Jahn O, Ganchev TD, Lhano MG, Marques MI. 2018. Tegmina-size variation in a Neotropical cricket with implications on spectral song properties. *Journal of Natural History* **52**:1225–1241. doi:10.1080/00222933.2018.1457728

Love RE, Walker TJ. 1979. Systematics and Acoustic Behavior of Scaly Crickets (Orthoptera: Gryllidae: Mogoplistinae) of Eastern United States. *Transactions of the American Entomological Society (1890-)* **105**:1–66.

Martins L de P, Redü DR, Oliveira GL de, Zefa E. 2012. Recognition characters and new records of two species of Phylloscyrtini (Orthoptera, Gryllidae, Trigonidiinae) from southern Brazil. *Iheringia, Sér Zool* **102**:95–98. doi:10.1590/S0073-47212012000100013

Martins LDP, Silva LGD, Henriques AL, Zefa E. 2013. First record of the genera Luzarida Hebard, 1928 and Luzaridella Desutter-Grandcolas, 1992 (Orthoptera, Gryllidae, Phalangopsinae) from Brazil, including a new species and description of the female of Luzarida lata Gorochov, 2011. *Zootaxa* **3609**:421–430. doi:10.11646/zootaxa.3609.4.4

McIntyre ME. 1977. Acoustical communication in the field crickets Pteronemobius nigrovus and P. bigelowi (Orthoptera: Gryllidae). *New Zealand Journal of Zoology* **4**:63–72. doi:10.1080/03014223.1977.9517937

Mello F de AG de, Pellegatti-Franco F. 1998. A New Cave Cricket of the Genus Endecous from Southeastern Brazil and Characterization of Male and Female Genitalia of E. itatibensis Rehn, 1918 (Orthoptera: Grylloidea: Phalangopsidae: Luzarinae). *Journal of Orthoptera Research* 185–188. doi:10.2307/3503517

Mello F de AG de, Reis JC dos. 1994. Substrate Drumming and Wing Stridulation Performed during Courtship by a New Brazilian Cricket (Orthoptera: Grylloidea: Phalangopsidae). *Journal of Orthoptera Research* 21–24. doi:10.2307/3503603

Metrani S, Balakrishnan R. 2005. The utility of song and morphological characters in delineating species boundaries among sympatric tree crickets of the genus Oecanthus (Orthoptera: Gryllidae: Oecanthinae): a numerical taxonomic approach. *orth* **14**:1–16. doi:10.1665/1082-6467(2005)14[1:TUOSAM]2.0.CO;2

Milach EM, Costa MKMD, Martins LDP, Nunes LA, Silva DSM, Garcia FRM, Oliveira ECD, Zefa E. 2016. New species of tree cricket Oecanthus Serville, 1831 (Orthoptera: Gryllidae: Oecanthinae) from Reserva Natural Vale, Espírito Santo, Brazil, with chromosome complement. *Zootaxa* **4173**:137–146. doi:10.11646/zootaxa.4173.2.4

Milach EM, Martins LDP, Costa MKMD, Gottschalk MS, Oliveira GLD, Redü DR, Neutzling AS, Dornelles JEF, Vasconcellos LA, Zefa E. 2015. A new species of tree crickets Oecanthus (Orthoptera, Gryllidae, Oecanthinae) in tobacco plantation from Southern Brazil, with body color variation. *Zootaxa* **4018**:266–278. doi:10.11646/zootaxa.4018.2.6

Nischk F, Otte D. 2000. Bioacoustics, Ecology and Systematics of Ecuadorian Rainforest Crickets (Orthoptera: Gryllidae: Phalangopsinae), with a Description of Four New Genera and Ten New Species. *Journal of Orthoptera Research* 229–254. doi:10.2307/3503651

Orsini MP, Costa MKMD, Szinwelski N, Martins LDP, Corrêa RC, Timm VF, Zefa E. 2017. A new species of Miogryllus Saussure, 1877 and new record of Miogryllus piracicabensis Piza, 1960 (Orthoptera: Gryllidae) from State of Rio Grande do Sul, Brazil, with calling song and chromosome complement. *Zootaxa* **4291**:361–372. doi:10.11646/zootaxa.4291.2.8

Otte D. 2007. Australian Crickets (Orthoptera: Gryllidae). Academy of Natural Sciences.

Pereira MR, Miyoshi AR, Martins LDP, Fernandes ML, Sperber CF, Mesa A. 2013. New Neotropical species of Hygronemobius Hebard, 1913 (Orthoptera: Grylloidea: Nemobiinae), including a brief discussion of male genitalia morphology and preliminary biogeographic considerations of the genus. *Zootaxa* **3641**:1–20. doi:10.11646/zootaxa.3641.1.1

Robillard T, Desutter-Grandcolas L. 2004. High-frequency calling in Eneopterinae crickets (Orthoptera, Grylloidea, Eneopteridae): adaptive radiation revealed by phylogenetic analysis. *Biological Journal of the Linnean Society* **83**:577–584. doi:10.1111/j.1095-8312.2004.00417.x

Robillard T, Ichikawa A. 2009. Redescription of Two Cardiodactylus Species (Orthoptera, Grylloidea, Eneopterinae): The Supposedly Well-Known C. novaeguineae (Haan, 1842), and the Semi-Forgotten C. guttulus (Matsumura, 1913) from Japan. *jzoo* **26**:878–891. doi:10.2108/zsj.26.878

Robillard T, Nattier R, Desutter-Grandcolas L. 2010. New species of the New Caledonian endemic genus Agnotecous (Orthoptera, Grylloidea, Eneopterinae, Lebinthini). *Zootaxa* **2559**:17–35. doi:10.11646/zootaxa.2559.1.2

Robillard T, Su YN. 2018. New lineages of Lebinthini from Australia (Orthoptera: Gryllidae: Eneopterinae). *Zootaxa* **4392**:241–266. doi:10.11646/zootaxa.4392.2.2

Robillard T, Tan MK. 2013. A taxonomic review of common but little knwon crickets from Singapore and the Philippines (Insecta: Orthoptera: Eneoptera: Eneopterinae) 21.

Robillard T, Yap S, Yngente MV. 2013. Systematics of cryptic species of Lebinthus crickets in Mount Makiling (Grylloidea, Eneopterinae). *Zootaxa* **3693**:49–63. doi:10.11646/zootaxa.3693.1.3

Souza-Dias PGB, Szinwelski N, Fianco M, Oliveira ECD, Mello FD a. GD, Zefa E. 2017. New species of Endecous (Grylloidea, Phalangopsidae, Luzarinae) from the Iguaçu National Park (Brazil), including bioacoustics, cytogenetic and distribution data. *Zootaxa* **4237**:454–470. doi:10.11646/zootaxa.4237.3.2

Su YN, Rentz DCF. 2000. Australian Nemobiine Crickets: Behavioral Observations and New Species of Bobilla Otte & Alexander (Orthoptera: Gryllidae: Nemobiinae). *Journal of Orthoptera Research* 5–20. doi:10.2307/3503626

Symes LB, Collins N. 2013. Oecanthus Texensis: A New Species of Tree Cricket from the Western United States. *orth* **22**:87–91. doi:10.1665/034.022.0203

Tan MK, Japir R, Chung AYC, Wahab RBHA. 2020. New taxa of crickets (Orthoptera: Grylloidea: Phaloriinae, Phalangopsinae, Itarinae and Podoscirtinae) from Borneo (Brunei Darussalam and Sandakan). *Zootaxa* **4810**:244–270. doi:10.11646/zootaxa.4810.2.2

Tan MK, Kamaruddin KN. 2016. A new species of Gryllotalpa mole cricket (Orthoptera: Gryllotalpidae: (Gryllotalpinae) from Peninsular Malaysia. *Zootaxa* **4066**:552–560. doi:10.11646/zootaxa.4066.5.3

Vicente NM, Olivero P, Lafond A, Dong J, Robillard T. 2015. Gnominthus gen. nov., a new genus of crickets endemic to Papua New Guinea with novel acoustic and behavioral diversity (Insecta, Orthoptera, Gryllidae, Eneopterinae). *Zoologischer Anzeiger - A Journal of Comparative Zoology* **258**:82–91. doi:10.1016/j.jcz.2015.06.005

Weissman DB, Gray DA. 2019. Crickets of the genus Gryllus in the United States (Orthoptera: Gryllidae: Gryllinae). *Zootaxa* **4705**:1–277. doi:10.11646/zootaxa.4705.1.1

Zefa E, Acosta RC, Timm VF, Szinwelski N, Marinho MAT, Costa MKMD. 2018. The Tree Cricket Neoxabea brevipes Rehn, 1913 (Orthoptera: Gryllidae: Oecanthinae) from the Brazilian southern Atlantic Forest: morphology, bioacoustics and cytogenetics. *Zootaxa* **4531**:554–566. doi:10.11646/zootaxa.4531.4.6

Zefa E, Neutzling AS, Redü DR, Oliveira GLD, Martins LDP. 2012. A new species of Oecanthus and Oecanthus lineolatus Saussure, 1897 from Southern Brazil: species description, including phallic sclerites, metanotal glands and calling song (Orthoptera: Gryllidae: Oecanthinae). *Zootaxa* **3360**:53–67. doi:10.11646/zootaxa.3360.1.2
